# Single-cell exploration of ovarian aging across mammalian models

**DOI:** 10.1101/2025.10.15.682639

**Authors:** Minhoo Kim, Justin Wang, Xucheng Liang, Ella Schwab, Bérénice A. Benayoun

## Abstract

Female reproductive lifespan is limited by a finite ovarian reserve, and its depletion leads to menopause and chronic disease risk. However, the molecular drivers of ovarian aging remain poorly understood, and preclinical models are not well benchmarked against human ovarian biology. We harmonized public single-cell and single-nucleus ovarian RNA-seq datasets from human, macaque, mouse, and goat for cross-species analysis of ovarian aging. Granulosa cell loss was the most consistent compositional change, and granulosa and theca cells showed the largest age-associated transcriptional shifts. Aging granulosa cells downregulated DNA repair genes and induced cell-cycle inhibitor genes across species, and shifted along an age-associated pseudotime trajectory in human and mouse. Using ASGARD, we nominated drug repurposing candidates with cross-species support that could reverse age-related transcriptional changes. Together, our meta-analysis identifies conserved and species-specific features of ovarian aging, which provides a framework to nominate potential interventions prolonging fertility or protecting against its systemic impacts.

## Introduction

Ovarian aging represents a fundamental biological process with profound implications for female reproductive capacity and systemic health ^1–3^. The ovary is characterized by a finite pool of oocytes established before birth, which progressively declines with age ^4–6^. This decline culminates in reproductive senescence and, in humans, menopause ^4–6^. The menopausal transition is not only characterized by the loss of fertility but also by systemic changes driven by the sharp reduction in sex-steroid production ^7^. These endocrine shifts contribute to accelerated frailty and increased susceptibility to age-associated diseases such as osteoporosis, cardiovascular disease, and neurodegeneration ^8,9^. With societal trends toward delayed childbearing, the consequences of ovarian aging are becoming increasingly pressing, both for individual health and for public health at large ^10^. Despite its significance, ovarian aging remains relatively underexplored at the molecular and cellular levels, and the mechanisms by which it shapes reproductive and systemic aging are not fully understood.

The ovary is a complex organ composed of oocytes, granulosa cells, theca cells, stromal cells, vascular cells, and immune populations ^11^. These cells form tightly regulated networks that support folliculogenesis, cyclic hormone production, and reproductive success ^12^. Single-cell and single-nucleus RNA sequencing (sc/snRNA-seq) are therefore powerful tools for capturing cell-type-specific changes and age-related shifts in ovarian heterogeneity. Over the past decade, sc/snRNA-seq datasets of ovaries from humans and animal models have been generated, spanning developmental to post-reproductive stages ^13–21^. However, these datasets are often analyzed independently with divergent pipelines, limiting their potential to yield comprehensive insights. An integrative meta-analysis of datasets generated across labs, species, and reproductive states is therefore an important step toward a unified roadmap of mammalian ovarian aging.

Here, we conduct a meta-analysis of publicly available ovarian sc/snRNA-seq datasets from humans and three mammalian model species ^14–20^. We harmonized datasets through standardized preprocessing, so as to establish consistent annotations and reduce technical variation across datasets. We then leveraged our harmonized datasets to quantify age-related shifts in ovarian cell-type proportions, and identified the populations most affected by aging. Next, to investigate molecular mechanisms most impacted by ovarian aging across cell types, we identified genes with consistent age-associated changes across species, performed pathway analyses, characterized co-expression modules associated with ovarian aging, and examined age-associated progression of granulosa cell states using pseudotime analysis. We also assessed therapeutic potential of ovarian aging signatures by linking age-associated transcriptional signatures to drug perturbation profiles to nominate compounds that might counteract molecular aspects of ovarian aging. By combining ovarian sc/snRNA-seq data across species and applying multiple analytic approaches, our study outlines both conserved and species-specific features of ovarian aging. Although limited by the number of datasets currently available, our findings suggest candidate biomarkers and pathways for further study and contribute to ongoing efforts to understand the biology of ovarian aging.

## Results

### Assembly of cross-species transcriptomic datasets for integrative ovarian aging analysis

To systematically investigate ovarian aging across vertebrates, we curated single-cell and single-nucleus transcriptomic datasets spanning humans, macaques, mice, and goats (Fig. 1a,b, Table S1). Human datasets collectively covered women from early adulthood (∼20s) through peri-menopause (∼late 40s) and post-menopause (∼50s): GSE202601 ^14^, GSE255690 ^15^ and Tabula Sapiens (TS) ^22^ datasets. Cynomolgus macaque ovaries were represented at reproductive (5 years) and aged (19 years), approaching the reported age of menopause (∼20 years): GSE130664 ^16^. Mouse datasets together captured ovarian aging from young adulthood (2–6 months) through peri-estropause (9–15 months) and post-estropause (>15 months): GSE290742 ^23^, GSE232309 ^17^, PRJNA863443 [AC, F+ and Veh] ^19^, E-MTAB-12889 and E-MTAB-11491 datasets ^18^. To note, for the PRJNA863443 dataset, we analyzed the three control sub-cohorts: natural aging (“AC”), wild-type controls from the *Foxl2* haploinsufficiency model (labeled “F+”), and vehicle-treated controls from the VCD chemically induced premature ovarian aging model (“Veh”) ^19^. Finally, we incorporated goat dataset PRJNA1010653, comparing young ovaries (2 years) to reproductively senescent (10 years) ones ^20^, though we note that it is limited to one sample per age group and is thus best be considered through the lens of cross-dataset/species comparisons.

**Fig. 1.**
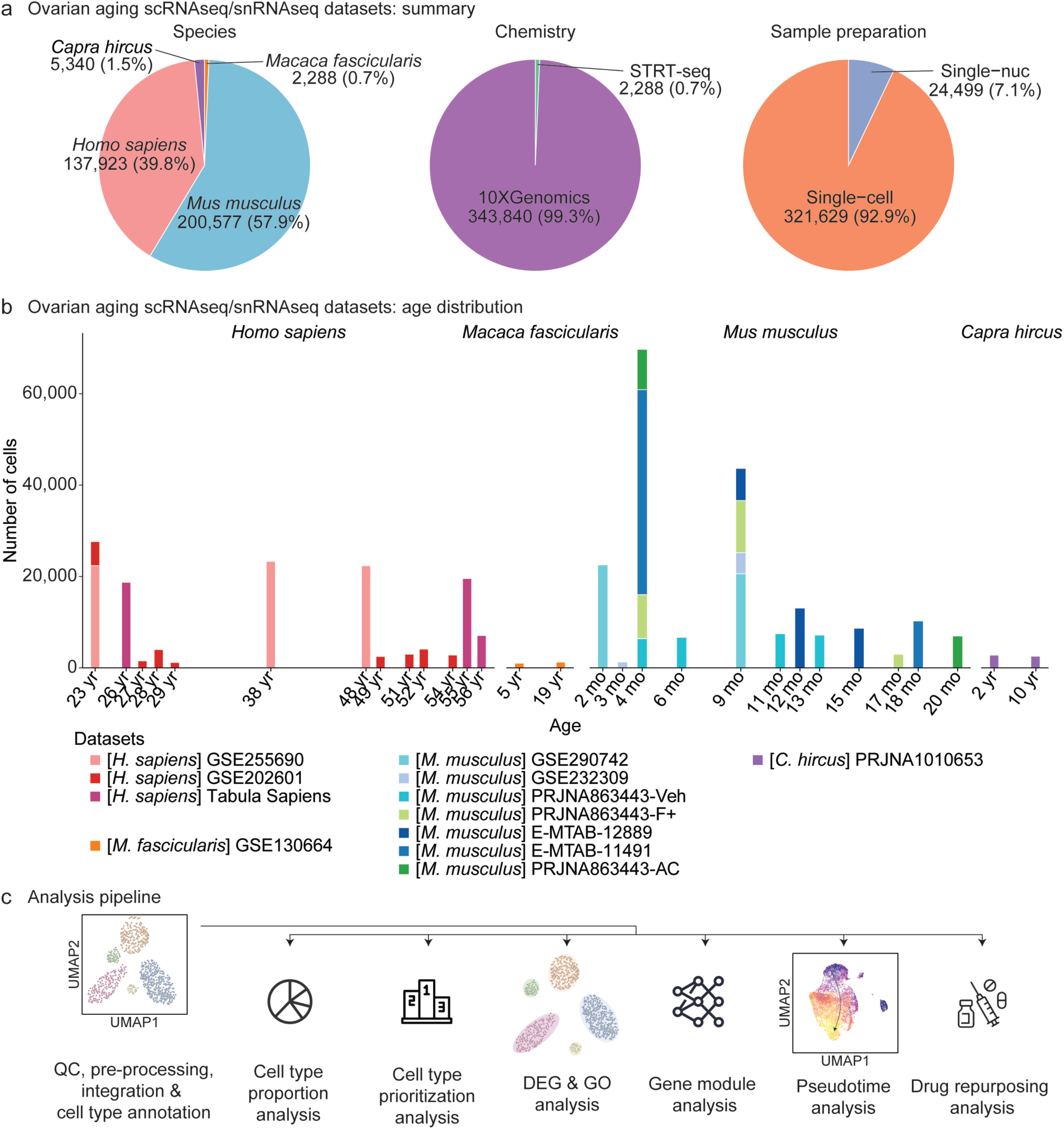
Cross-species datasets and integrated analysis framework for ovarian aging. (**a**) Summary of curated sc/snRNA-seq datasets. (**b**) Age coverage for each dataset. (**c**) Scheme for integrative analysis of ovarian aging single-cell and single-nucleus datasets across species.

These datasets were contextualized relative to species-specific reproductive milestones (Supplementary Fig. S1). In humans, perimenopause typically begins around 45 years, with menopause occurring at ∼52 years ^24^. In cynomolgus macaques, post-menopause arises around 20 years, though perimenopause is less well defined ^25^. In mice, estropause is used in place of menopause ^26^. Peri-estropause is commonly defined as 9–12 months of age ^27^, but the transition to acyclicity has been reported to extend to approximately 15 months ^26^. We therefore defined peri-estropause as 9–15 months and post-estropause as >15 months. In goats, reproductive aging is marked by reproductive senescence at approximately 6 years, but a discrete perimenopause-like interval has not been clearly defined ^28^. This alignment provided a framework for comparing age-associated features across species despite differences in reproductive biology.

To enable integrative analysis, each dataset was uniformly pre-processed, quality-controlled and annotated (Fig. 1c). We then examined age-associated changes at multiple levels: (i) shifts in cell type composition, (ii) global transcriptional perturbations and cell type prioritization using Augur ^29^, and (iii) pseudobulk-level differential gene expression across key cell types, identifying genes with consistent age-associated changes across species to identify conserved signatures. We explored functional implications through gene ontology enrichment and weighted gene co-expression network analysis (WGCNA) ^30^, which identified modules associated with ovarian functional decline. Importantly, to examine age-associated and maturation-associated changes along the transcriptional trajectory of granulosa cells (the key steroidogenic cell type of the mammalian ovary), we performed pseudotime analysis using Monocle3 ^31^. Finally, we applied ASGARD ^32^ to nominate compounds predicted to reverse ovarian aging-associated transcriptional signatures, revealing candidate ovarian gerothepeutics. Together, this framework delineates conserved and species-specific features of ovarian aging at single-cell resolution.

### Identification of major ovarian cell types across species

To establish a consistent cellular framework for cross-species comparisons, we performed unsupervised clustering and visualization of ovarian single-cell and single-nucleus transcriptomic profiles using UMAP separately for each species (Fig. 2). To note, for humans and mice, where multiple datasets were available, we first integrated all datasets within the specific species to account for technical variation before embedding. This step was not needed for macaques and goats, where only one dataset per species was available.

**Fig. 2.**
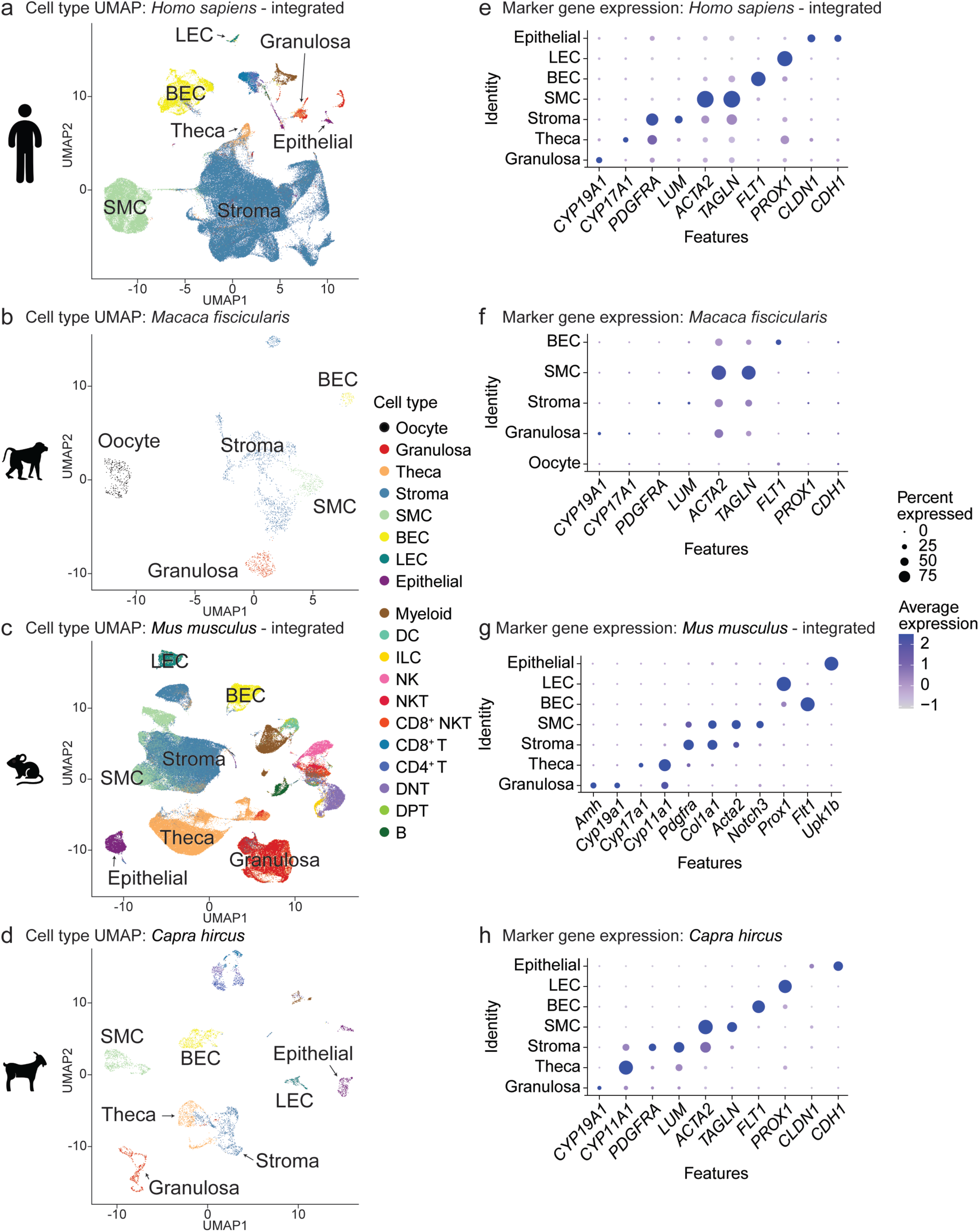
Cross-species cell-type landscape: UMAPs and marker gene profiles. UMAP plots of integrated human (**a**), macaque (**b**), integrated mouse (**c**) and goat (**d**) datasets. Marker gene dot plots of *Ptprc^-^* cell types from integrated human (**e**), macaque (**f**), integrated mouse (**g**) and goat (**h**) datasets. SMC: smooth muscle cells; BEC: blood endothelial cells; LEC: lymphatic endothelial cells; DNT: double-negative T cells; DPT: double-positive T cells.

Across species, we were able to consistently identify key ovarian cell types, including granulosa cells, theca cells, and stromal cells (Fig. 2b-d). In addition, multiple immune populations were detected, including myeloid cells as well as subsets of T and B cells, reflecting the known immunological complexity of the ovarian microenvironment (Supplementary Fig. S2). By contrast, detection of oocytes was extremely limited, most likely due to technical constraints of droplet-based single-cell platforms such as 10x Genomics, which exclude very large cell types during library preparation to prevent microfluidics clogging. Together, our analyses robustly identify a conserved set of major ovarian cell populations across mammals, providing a reference for subsequent interrogation of aging-associated cellular and molecular changes. To facilitate community access and reuse, we developed an interactive R Shiny application that allows users to explore our processed single-cell and single-nucleus ovarian datasets across species (https://minhooki.shinyapps.io/cross-species-ovary-scRNA/).

### Age-associated shifts in ovarian cell-type composition across species

We next quantified age-associated changes in cell-type proportions using scProportionTest ^33^, which estimates permutation-based p-values per cluster and returns bootstrap confidence intervals for the magnitude of change (Fig. 3a). To note, since the macaque dataset was generated by hand-picking cells for STRT-seq, its sampling fractions do not reflect the original tissue composition, so we excluded it from proportion analyses to avoid this confounding factor. Because sampled ages varied across datasets (Supplementary Fig. S1a), samples were grouped by reproductive phase and the youngest and oldest phases were compared. For datasets spanning all three phases or a single phase, we adjusted the comparison accordingly (see Methods).

**Fig. 3.**
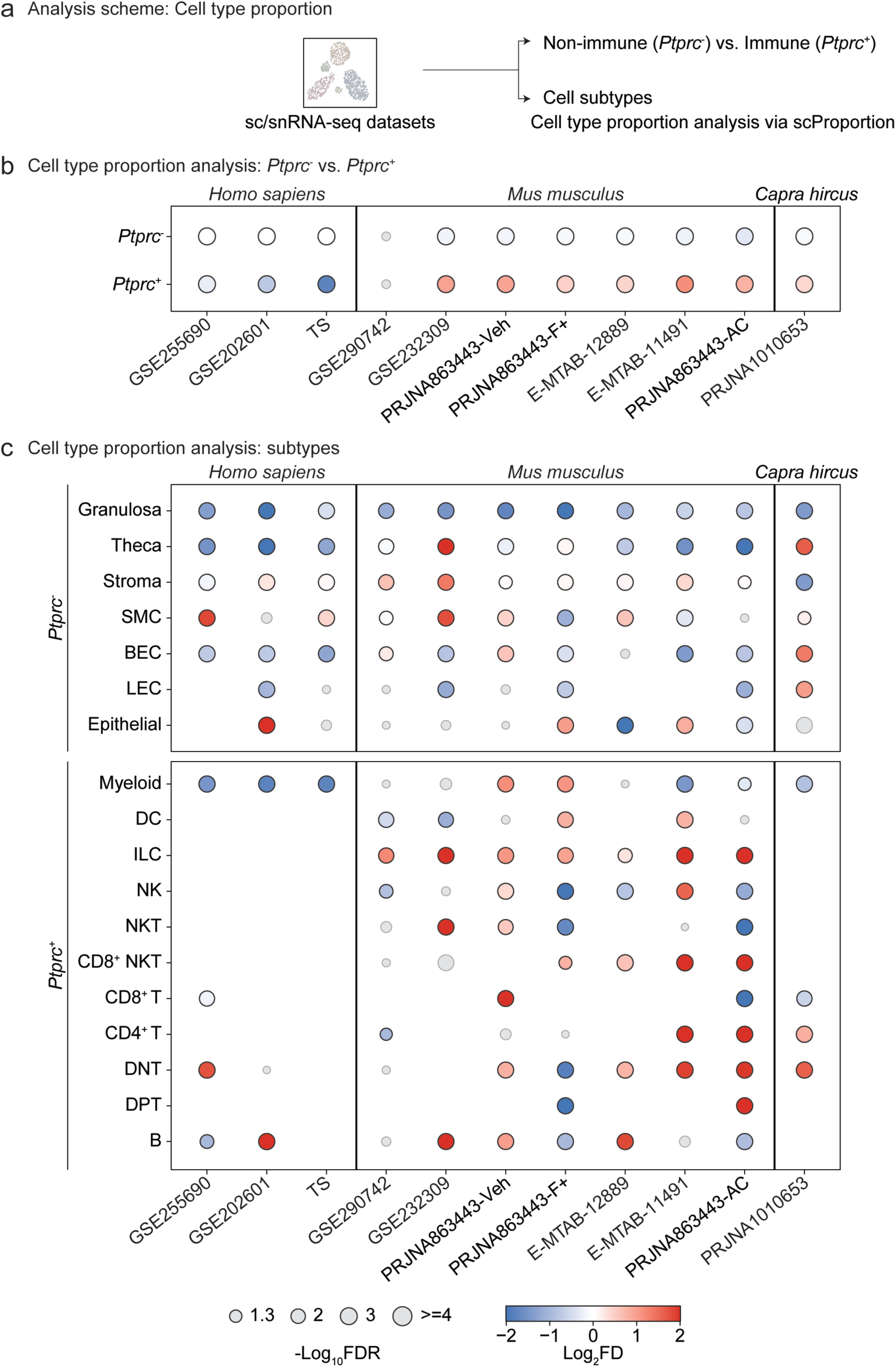
Analysis of ovarian aging-associated cell type proportion changes across datasets. (**a**) Schematic of the analysis workflow. (**b**) Proportion differences of *Ptprc*^-^ and *Ptprc*^+^ cells in human, mouse, and goat datasets. (**c**) Proportion differences of cell subtypes detected in human, mouse, and goat datasets. Significant comparisons (FDR < 0.05) are shown in color; non-significant comparisons are shown in gray. SMC: smooth muscle cells; BEC: blood endothelial cells; LEC: lymphatic endothelial cells; DNT: double-negative T cells; DPT: double-positive T cells.

We first analyzed shifts using a broad partition of immune (*Ptprc*⁺) versus non-immune (*Ptprc*⁻) cells (Fig. 3b). Interestingly, while immune-cell proportions decreased with age in all three human datasets, they increased in six of seven mouse datasets and in goat. At the level of cell-type subpopulations (Fig. 3c), granulosa cells decreased with age in every dataset across all three species analyzed (including humans), representing the most concordant trend. Theca cells declined in all human datasets and in many mouse datasets (e.g., E-MTAB-11491 and PRJNA863443-AC), but increased in GSE232309 and goat. Stromal cells showed no predominant direction. Within the immune compartment, myeloid cells decreased in all human datasets, in line with the reduced immune fraction, whereas lymphoid populations expanded in model organisms. Innate lymphoid cells (ILCs) increased in every mouse dataset. Increases in double-negative T (DNT), CD4⁺ T, and CD8⁺ NKT cells were most pronounced in the comparisons extending to the oldest ages (E-MTAB-11491 and PRJNA863443-AC, 18 and 20 months). DNT and CD4⁺ T cells also increased in goat.

While human dataset-specific observations may be due to true species-specificity, a combination of both biological and technical factors could also explain such discrepancies which are worth noting: (i) human data is derived from small ovarian cortical biopsies, since this is where most of ovarian follicles reside (in contrast to the more vascularized medulla), (ii) mouse and goat samples were obtained from the dissociation of the entire tissue, including cortical and medullar regions. Thus, because age-associated immune expansion in the ovary can be spatially concentrated, particularly within corpora lutea and atretic follicles in aged mice ^34^, the immune fraction captured in a given sample depends on the structures it contains. Human cortical biopsies therefore cannot be directly compared with whole-ovary mouse and goat datasets in terms of cell composition.

Together, these analyses identify granulosa-cell loss as a conserved feature of ovarian aging, whereas immune remodeling is not consistent across datasets. This divergence warrants follow-up using harmonized sampling and future validation studies taking into account the unique and shared histological features of ovarian organization across mammalian species.

### Cell-type prioritization highlights conserved transcriptional perturbations in granulosa and theca cells

We applied Augur ^29^ to quantify the strength of ovarian aging-associated transcriptional perturbations within each cell type (Fig. 4a). For cross-species comparison, we restricted to cell types detected in 2 or more species and aggregated within-dataset ranks via a fractional rank product (see Methods). Among cell types represented in 2 or more species, granulosa and theca consistently occupied the top ranks, indicating the strongest, most widespread aging-associated shifts (Fig. 4b; Supplementary Fig. S3).

**Fig. 4.**
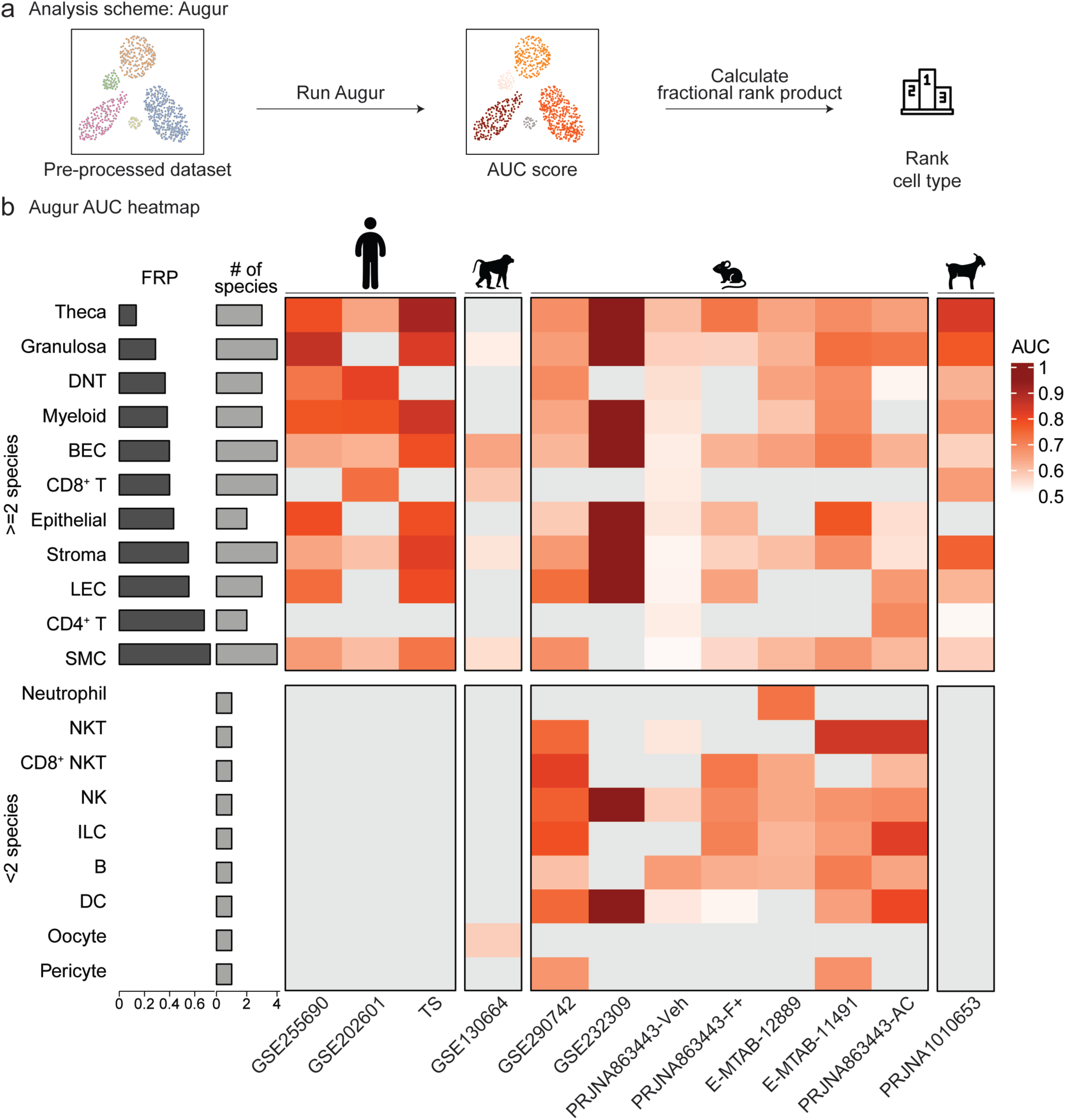
Analysis of ovarian aging–associated cell type transcriptional perturbations across datasets using Augur. (**a**) Schematic of global gene expression analysis comparisons using Augur. (b) Heatmap of Augur AUC scores, ordered by fractional rank product, for cell types present in at least three datasets. SMC: smooth muscle cells; BEC: blood endothelial cells; LEC: lymphatic endothelial cells; DNT: double-negative T cells; DPT: double-positive T cells.

Immune subsets tended to show intermediate or species-dependent ranks, and stromal compartments were variable across datasets. Guided by these results and biological relevance, we focused downstream analyses on granulosa, theca, and stroma: granulosa and theca constitute the steroidogenic follicular unit governing ovarian estrogen/progesterone production ^35,36^, while stroma comprises a large fraction of the ovary and shapes the extracellular matrix, vasculature, and paracrine milieu ^37^.

### Gene-level analysis identifies cross-species, age-associated signatures in granulosa, theca, and stromal cells

To identify the genes underlying these cell-type-level perturbations, we performed pseudobulk differential expression analysis (Fig. 5a; see Methods). Within each dataset and cell type, counts were aggregated by sample, and age contrasts were tested with DESeq2 ^38^. Granulosa, theca, and stromal cells showed the highest fraction of significant DEGs (FDR < 0.05; Fig. 5b). To identify conserved signatures, gene IDs were humanized using orthology mapping and p-values were combined within species using metaRNASeq ^39^. For granulosa and stromal cells, we then retained genes that were significant (FDR < 10%) with the same direction of change in at least 3/4 species. For theca, which was not detected in the macaque dataset, genes had to meet this criterion in all 3 species. Intriguingly, conserved signals were shared mainly by human, mouse, and goat, with the macaque contributed few overlapping genes, which may be a technical byproduct of the relative sparsity of the macaque dataset (Fig. 5c, d; Supplementary Figs. S4-S6).

**Fig. 5.**
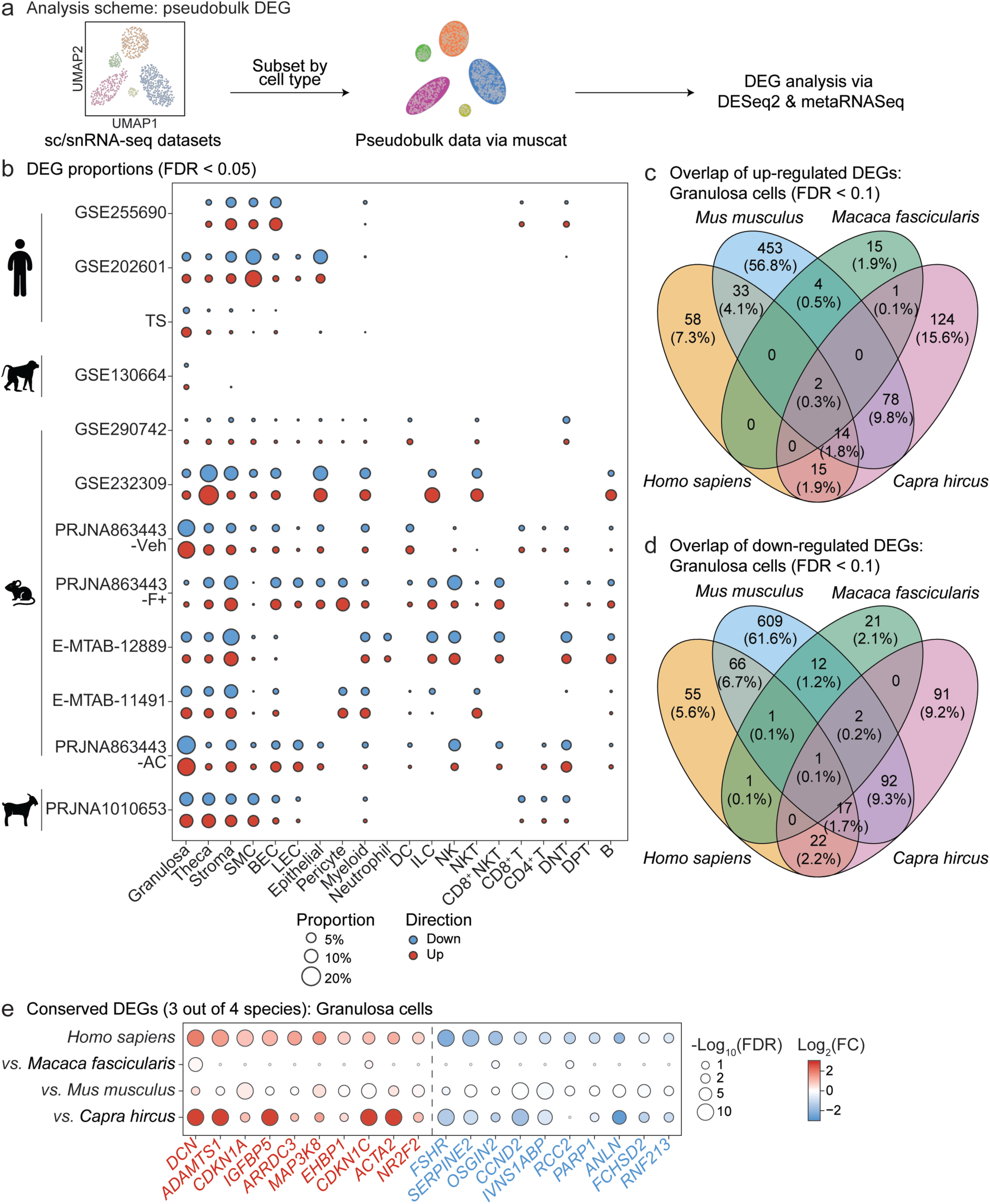
Analysis of ovarian aging–associated differential gene expression in granulosa cells across datasets. (**a**) Schematic of pseudobulk analysis pipeline. (**b**) Bubble plot of significant DEG proportions (FDR < 0.05) by cell type across datasets. (**c**) Venn diagram of overlapping up-regulated DEGs (FDR < 0.1) across species for granulosa cells. (**d**) Venn diagram of overlapping down-regulated DEGs (FDR < 0.1) across species for granulosa cells. (**e**) Bubble plot of top conserved DEGs (3 out of 4 species; FDR < 0.1) for granulosa cells. SMC: smooth muscle cells; BEC: blood endothelial cells; LEC: lymphatic endothelial cells; DNT: double-negative T cells; DPT: double-positive T cells.

In granulosa cells, 16 upregulated and 17 downregulated genes met the conservation criterion (Fig. 5c–e; Supplementary Fig. S4a). Upregulated genes included the cyclin-dependent kinase inhibitors *CDKN1A* and *CDKN1C* ^40^ and the stress-responsive genes *BTG2* and *TXNIP* ^41,42^. Downregulated genes included *FSHR* and *CCND2*, which mediate FSH-driven granulosa cell proliferation ^43,44^, the DNA repair genes *BRCA1* and *PARP1* ^45,46^, and the proliferation-associated genes *ANLN* and *RCC2* ^47,48^. This pattern is consistent with reduced gonadotropin responsiveness and proliferative capacity in aged granulosa cells.

In theca cells, 9 upregulated and 6 downregulated genes were shared across human, mouse, and goat (Supplementary Fig. S5). Downregulated genes included *DHCR24* and *ACSL4* ^49,50^, which function in cholesterol synthesis, and lipid metabolism, respectively. *DCN* and *MAP3K8* ^51,52^ were upregulated in both theca and granulosa cells.

In stromal cells, 14 upregulated and 31 downregulated genes were shared across human, mouse, and goat (Supplementary Fig. S6). Downregulated genes were dominated by extracellular matrix components (*COL1A1*, *COL1A2*, *COL4A1*, *COL5A2*, *FN1*, *SPARC*, *CCN2*) ^53,54^, contractile genes (*ACTA2*, *DES*, *MYH11*, *TAGLN*, *MYLK*) ^55^, and proliferation-associated genes (*CENPE*, *KIF20B*, *UHRF1*) ^56^. Upregulated genes included the immunoproteasome components *PSMB9* and *PSME1* and the iron exporter *SLC40A1* ^57,58^.

Together, these results define conserved age-associated transcriptional changes in each major somatic compartment of the ovary: reduced FSH-responsive and proliferative programs in granulosa cells, altered cholesterol handling in theca cells, and reduced extracellular matrix and contractile gene expression in stromal cells.

### GO enrichment highlights conserved age-associated pathway shifts in granulosa and stromal cells

To place these gene-level changes in a pathway context, we first performed Gene Ontology (GO) enrichment of cell-type-resolved pseudobulk data using multi-contrast enrichment analysis with the mitch framework ^59^ (Fig. 6a; Supplementary Fig. S7a). In granulosa cells, terms related to autophagy, lysosome, late endosome membrane, and inflammatory response were elevated with age across most datasets. In contrast, terms related to genome maintenance, including DNA repair, chromosome telomeric region, and chromatin organization, were reduced, as was cell adhesion (Fig. 6b). We independently analyzed our data using Reactome gene sets as well, which showed concordant patterns: autophagy terms increased and non-integrin membrane-ECM interactions decreased in most datasets (Supplementary Fig. S7b). The reduced enrichment of DNA repair terms is consistent with the conserved downregulation of *BRCA1* and *PARP1* (Fig. 5e; Supplementary Fig. S4a). Cell cycle-related terms (e.g., APC/C-mediated degradation, synthesis of DNA), however, changed in varying directions across datasets. In theca cells, regulation of gene expression was modestly elevated in most datasets, but no robust pathway-level signature emerged (Fig. 6c; Supplementary Fig. S7c).

**Fig. 6.**
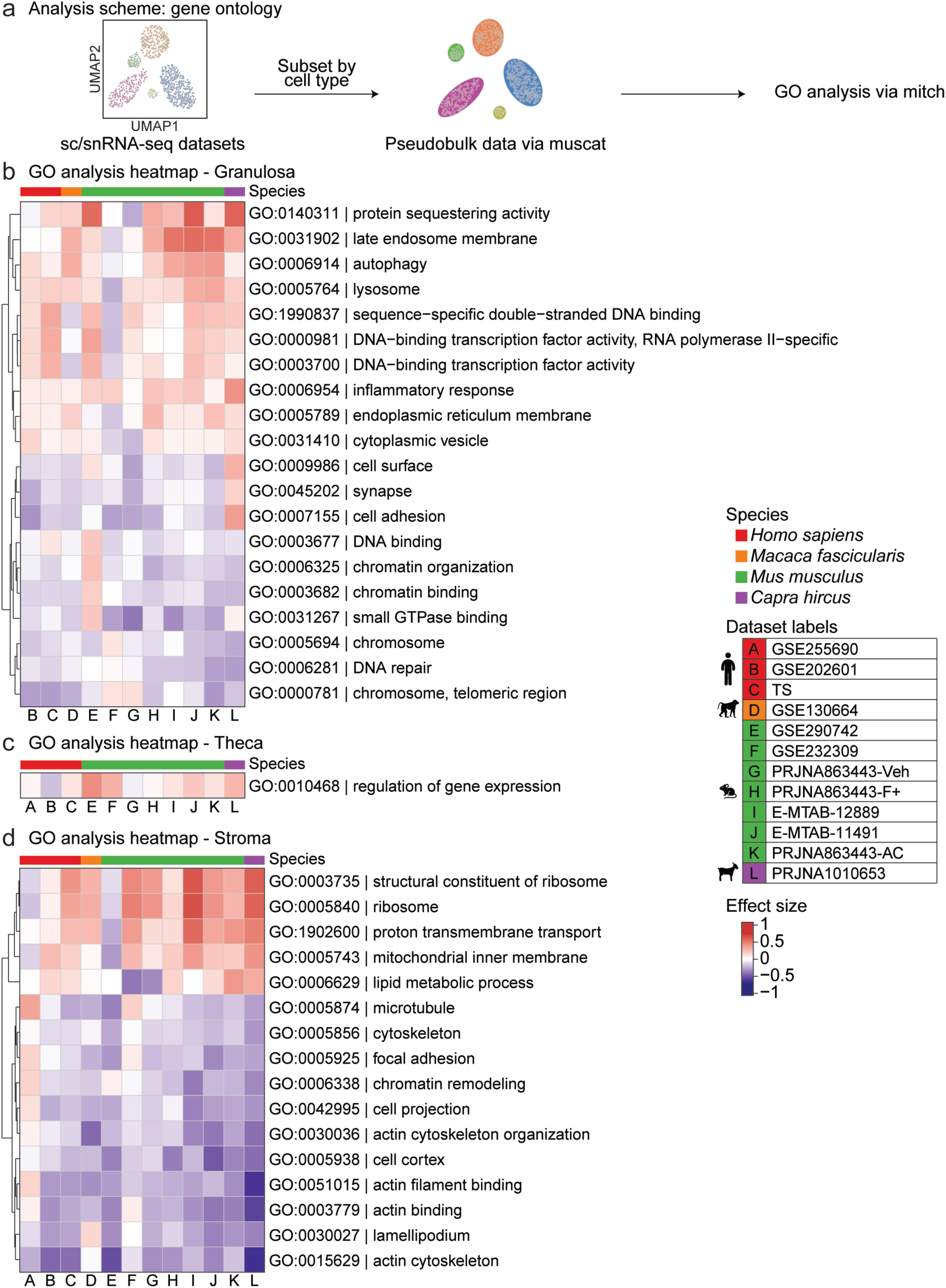
Analysis of ovarian aging–associated gene ontology terms across datasets using mitch. (**a**) Analysis scheme of gene ontology analysis for granulosa, theca and stromal cells across datasets. (**b-d**) Heatmap of GO terms detected (FDR < 0.05) in granulosa (**a**), theca (**b**) and stromal (**c**) cells.

In stromal cells, terms related to the ribosome, proton transmembrane transport, and mitochondrial inner membrane were elevated with age in most datasets (Fig. 6d). The corresponding Reactome terms (translation, mitochondrial translation, complex I biogenesis, and respiratory electron transport) showed the same pattern (Supplementary Fig. S7d). GSE255690 and GSE290742 were exceptions and showed reduced mitochondrial translation. Conversely, cytoskeletal and adhesion terms, including actin cytoskeleton, actin binding, focal adhesion, lamellipodium, and microtubule, decreased broadly, together with Rho GTPase signaling. This is consistent with the reduced expression of ECM and contractile genes among conserved stromal DEGs (Supplementary Fig. S6c).

Together, these results point to increased lysosomal, autophagic, and inflammatory programs and reduced genome maintenance in aging granulosa cells. Aging stromal cells showed increased translational and mitochondrial programs and reduced cytoskeletal and adhesion programs.

### Module-level signatures reveal a conserved age-repressed granulosa program across species

Next, WGCNA was used to identify co-expression programs associated with ovarian aging. Consensus networks were built from pseudobulk profiles of human datasets for granulosa (GSE202601 and TS), theca (GSE255690 and TS), and stromal cells (GSE202601, GSE255690 and TS) (Fig. 7a; Supplementary Figs. S8, S9). Modules were tested for association with age (FDR < 10%), and age-associated module gene sets were evaluated by GSEA in each dataset to test cross-species consistency (Fig. 7a). No WGCNA stromal module reached significance.

**Fig. 7.**
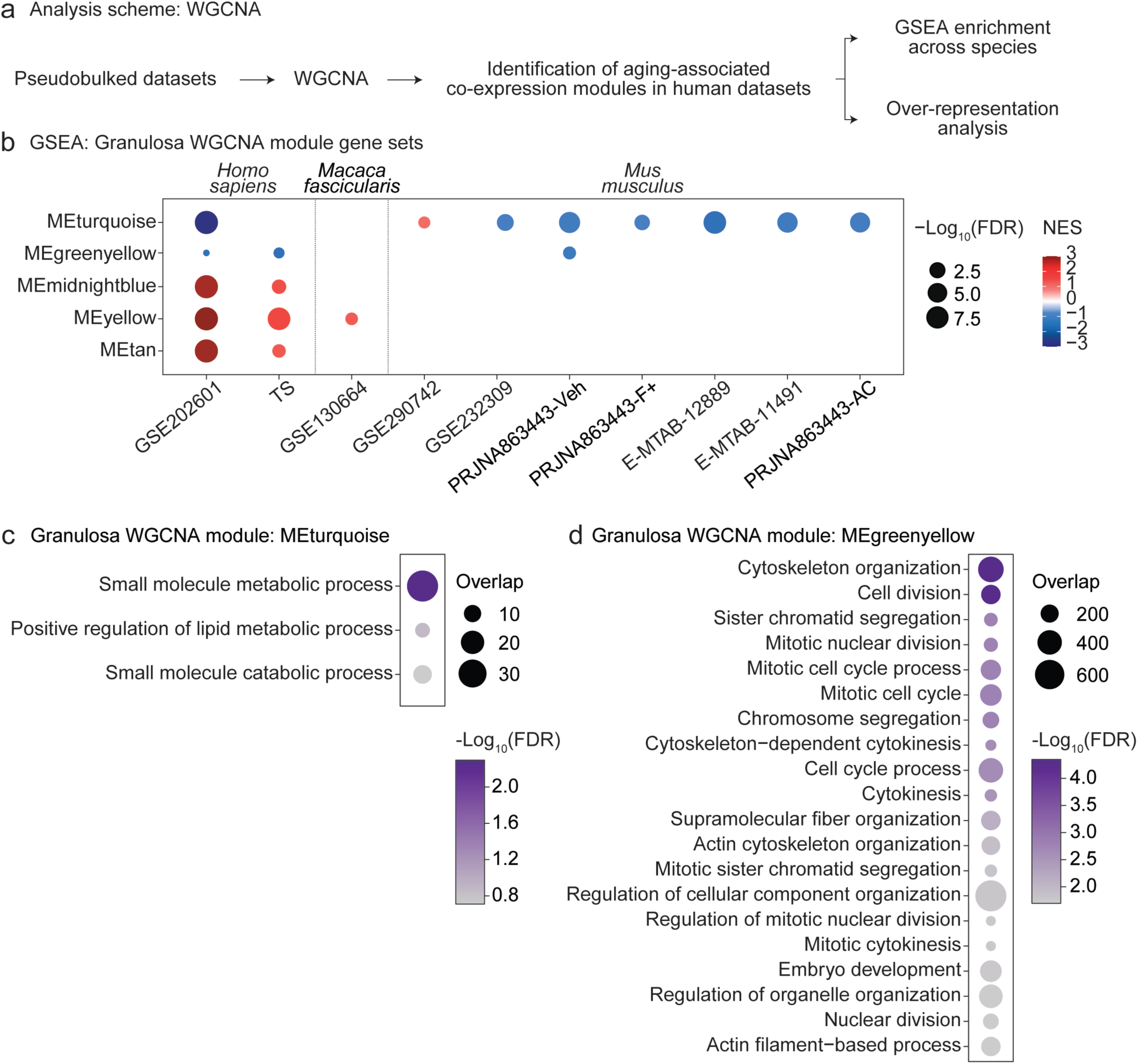
Analysis of ovarian aging–associated granulosa co-expression modules across datasets. (**a**) Schematic of WGCNA analysis framework. (**b**) Cross-species GSEA enrichment (FDR < 0.1) of human granulosa cells-derived WGCNA module gene sets against macaque, mouse and goat datasets. (**c, d**) Over-representation analysis of gene ontology terms for granulosa WGCNA modules (FDR < 0.1).

In granulosa cells, five modules were associated with age: two (turquoise and greenyellow) were negatively associated, and three (midnightblue, yellow, and tan) were positively associated (Supplementary Fig. S8b). GSEA recapitulated these directions in both human datasets (Fig. 7b). The turquoise module was the most conserved: it was depleted with age in six of seven mouse datasets. In contrast, the positively associated modules were largely not recapitulated outside human, with the exception of yellow in macaque. None of the module gene sets was significantly enriched in goat. ORA showed that the turquoise module was enriched for small molecule metabolic process (Fig. 7c). The greenyellow module was enriched for cell division, chromosome segregation, cytokinesis, and cytoskeleton organization (Fig. 7d), consistent with the conserved downregulation of proliferation-associated genes such as *CCND2*, *ANLN*, and *RCC2* (Fig. 5e).

In theca cells, two modules (turquoise and lightgreen) were negatively associated with age, and one (grey60) was positively associated (Supplementary Fig. S9b). These directions were consistent across both human datasets but varied in mouse (Supplementary Fig. S10a). For example, the turquoise module was depleted with age in GSE290742 and GSE232309 but enriched in three other mouse datasets. The turquoise module was strongly enriched for oxidative phosphorylation, mitochondrial gene expression, and translation (Supplementary Fig. S10b), indicating reduced mitochondrial and translational programs in aging human theca. Grey60 terms, including extracellular matrix organization and cholesterol transport, did not reach significance (Supplementary Fig. S10c).

Together, these analyses identify a granulosa metabolic module as the most conserved age-repressed program across human and mouse. Theca modules were consistent within human, but not across other species.

### Pseudotime analysis reveals age-associated progression of granulosa cell states in human and mouse

Because granulosa cells showed the most consistent age-associated changes across analyses, we next examined whether aging shifts granulosa cells along a defined transcriptional trajectory, using pseudotime analysis of granulosa cells integrated separately for human and mouse (Fig. 8a–c). Mean pseudotime per donor or animal differed across age groups in both species (Kruskal-Wallis p = 0.019 for human and p = 0.0058 for mouse; Fig. 8d, e). In human, pseudotime was higher in old (>50 years) than young (30–50 years) donors (adjusted p = 0.041). In mouse, pseudotime was higher in middle-aged (9–15 months) than young (<9 months) animals (adjusted p = 0.012), with a similar trend in old animals (>15 months; adjusted p = 0.063).

**Fig. 8.**
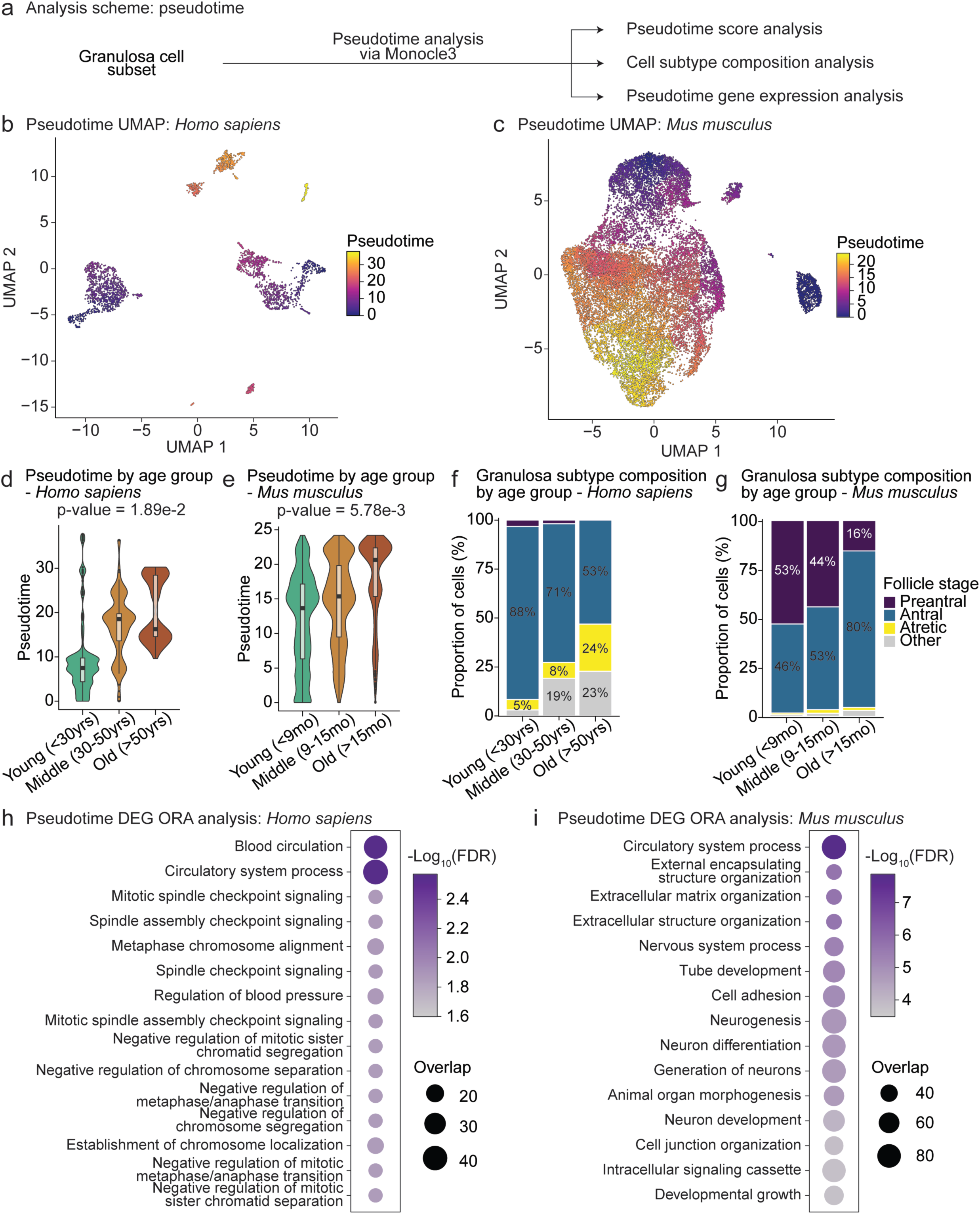
Granulosa cell states shift along an age-associated pseudotime trajectory in human and mouse. (**a**) Schematic of the pseudotime analysis workflow. (**b,c**) UMAP embeddings of integrated granulosa cells from (**b**) human and (**c**) mouse, colored by monocle3 pseudotime. (**d, e**) Pseudotime distributions by age group in (**d**) human, and (**e**). Violin plots show per-cell pseudotime, and embedded box plots indicate the median and interquartile range. Statistical testing was performed on median pseudotime per donor or animal (human, n = 14 donors; mouse, n = 52 animals) using the Kruskal-Wallis test, and p-values are shown above each panel. (**f**, **g**) Follicle-stage composition of granulosa cells (preantral, antral, atretic, and other) by age group in (**f**) human and (**g**) mouse, shown as the percentage of granulosa cells in each group. (**h**, **i**) Over-representation analysis of genes that change monotonically along pseudotime in (**h**) human and (**i**) mouse.

Aging was also accompanied by shifts in granulosa cell subtype composition, although the nature of the shift differed between species. In human, the proportion of antral granulosa cells declined with age (88%, 71%, and 53% in young, middle, and old groups), whereas the proportion of atretic granulosa cells increased (5%, 8%, and 24%) (Fig. 8f). In mouse, preantral granulosa cells declined (53%, 44%, and 16%) and antral granulosa cells increased (46%, 53%, and 80%), while atretic cells remained rare (Fig. 8g).

We next identified genes that changed monotonically along pseudotime (Supplementary Fig. S11). Pseudobulk expression of these genes largely followed chronological age, and increasing genes were most highly expressed in the oldest samples of each species (Supplementary Fig. S11a, b). ORA of the combined increasing and decreasing genes showed enrichment of spindle assembly checkpoint and chromosome segregation terms, as well as blood circulation, in human (Fig. 8h). In mouse, enriched terms included extracellular matrix organization, cell adhesion, and developmental processes such as neurogenesis and tube development (Fig. 8i). Circulatory system process was enriched in both species.

Together, these results define a granulosa cell trajectory that advances with age in both human and mouse. Along this trajectory, human granulosa cells shift toward atretic states, whereas mouse granulosa cells shift away from preantral states.

### ASGARD-based drug repurposing nominates cross-species–supported candidates

Finally, we asked if we could leverage the power of cross-species ovarian aging associated signatures to nominate potential interventions that may reverse aspects of transcriptional shifts with aging in the ovary. For this purpose, we applied ASGARD ^32^ and the LINCS1000 reference perturbation library to identify compounds predicted to reverse age-associated transcriptional changes in the ovary in each dataset, and results were combined within and across species to identify cross-species supported drug repurposing candidates (Fig. 9a). All ten top-ranked compounds reached significance in mouse, the species with the most datasets, whereas support in other species varied (Fig. 9b). Excitingly, dexamethasone showed the broadest support, reaching significance independently in human, macaque, and mouse.

**Fig. 9.**
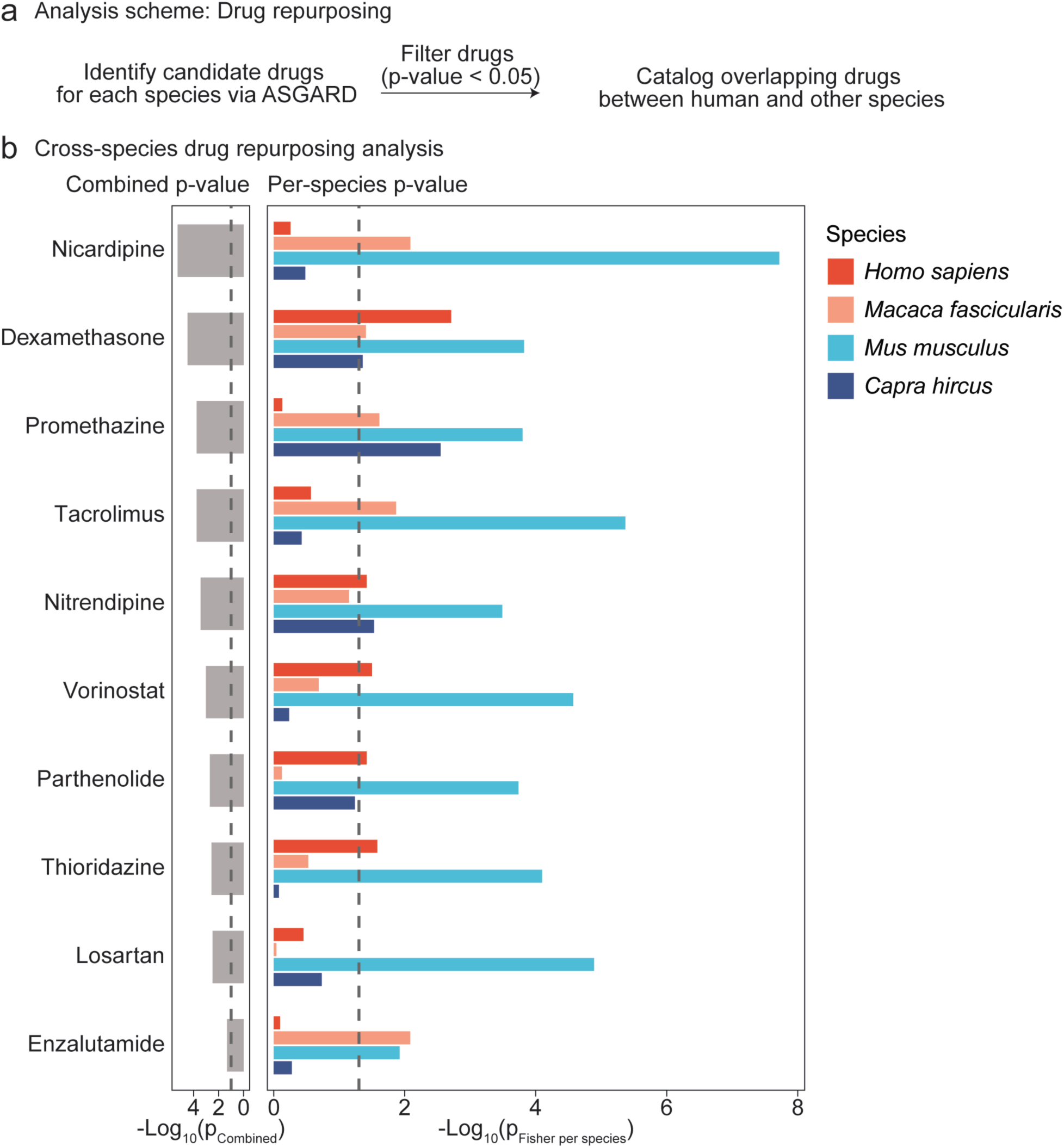
Drug-repurposing analysis for ovarian aging signatures across datasets using ASGARD. (**a**) Schematic of drug repurposing analysis framework. (**b**) Grouped bar plot of −log_10_(combined P) per compound and species, where combined P is computed within species using Fisher’s method to aggregate p-values. Bars are shown for the top 10 compounds significant in combined P.

Nitrendipine was significant in human, mouse, and goat, and vorinostat, parthenolide, and thioridazine were significant in human and mouse. The top-ranked compound, nicardipine, was supported by mouse and macaque, but not human data. Among the next-ranked compounds, ixazomib, tamoxifen, and troglitazone were significant in human, but had limited support in other species (Supplementary Fig. S12). These candidates warrant future mechanistic evaluation to determine whether they may be tractable candidate drugs for interventions to slow down ovarian aging. Larger age-stratified single-cell cohorts should increase power and cross-species concordance.

## Discussion

In this meta-analysis study, we found that, among ovarian cells, granulosa and theca cells showed the largest age-associated transcriptional shifts across species, and age-related decrease in granulosa cell abundance is the most consistent compositional change in the aging mammalian ovarian tissue. Several factors may underlie such larger shifts in these follicular somatic cells: cumulative proliferative and steroidogenic demand across cycles; increasing gonadotropin drive as follicles are depleted, which can promote receptor desensitization or drift in downstream signaling; microenvironmental remodeling (*e.g*., ECM stiffening, altered vascular tone) that constrains nutrient and oxygen delivery; and reduced chromatin regulatory capacity that limits rapid state transitions ^60,61^. These possibilities align with the conserved signatures we observed, including cross-species downregulation of *FSHR* and *CCND2*, induction of the cell-cycle inhibitors *CDKN1A* and *CDKN1C*, reduced DNA repair and chromatin terms, and an age-associated shift forward in granulosa pseudotime. Physiologically, such changes could contribute to attenuated hormone responsiveness, reduced estradiol and progesterone output, impaired ovulatory responses, and suboptimal luteal function, with downstream effects on cycle regularity and hypothalamic-pituitary-ovarian feedback ^62^.

Stromal cells showed conserved gene-level and pathway-level changes, but their proportions varied across datasets and no stromal co-expression module was associated with age. This pattern likely reflects the greater cellular and regional heterogeneity of the stromal compartment of the ovary (including potential fibroblast, perivascular, endothelial, and immune subsets) and variable cortical and medullary sampling across studies. Subtype-resolved annotation and region-aware sampling (or spatial transcriptomic analyses) will be crucial to better define stromal aging programs across species with higher confidence.

Immune remodeling diverged between species: immune cell proportions decreased with age in human datasets but increased in mouse and goat. The lymphoid expansion we observed in aged mouse ovaries, including ILCs and DNT cells, is consistent with previous single-cell studies ^17,63^. In humans, a recent single-nucleus atlas of postmenopausal ovaries likewise reported age-associated immune depletion, and showed that parity opposed many age-associated changes ^55^. However, we should note that spatial transcriptomics of the mouse ovary across the estrus cycle and with aging identified age-associated immune cell expansion to be primarily spatially restricted to corpora lutea and atretic follicles, including T cell expansion within aged corpora lutea ^34^. Thus, although the divergence between human and other species may reflect biology, it is also possible that inherent sampling strategy differences (e.g., cortical fragments versus whole organs), limited donor numbers, and individual factors (e.g., parity and hormone use) may drive the difference in the datasets. Harmonized sampling across species, with reproductive history recorded, will be needed to distinguish these possibilities.

Meta-analysis of pseudobulk DEGs identified genes with consistent age-associated changes in ≥3 of the analyzed species. In granulosa cells, conserved downregulation of *FSHR* and *CCND2* points to reduced FSH-driven signaling and cell proliferation. *FSHR* encodes the granulosa FSH receptor, which couples to cAMP/PKA to drive aromatase induction, LH receptor expression acquisition, and steroidogenesis ^64^, and *CCND2* is an FSH-responsive cyclin required for granulosa cell proliferation ^44^. The reduction in *CCND2*, together with induction of *CDKN1A* and *CDKN1C*, suggests a shift toward cell-cycle restraint in aged granulosa cells. Downregulation of *BRCA1* and *PARP1* extends previous links between impaired DNA double-strand break repair and ovarian aging, established mainly in oocytes ^45,46^, to the granulosa compartment.

In theca cells, downregulation of *DHCR24* and *ACSL4*, which function in cholesterol synthesis and lipid metabolism, suggests reduced capacity to supply substrate for thecal androgen synthesis ^49,50^. The aging ovarian stroma becomes increasingly fibrotic and stiff ^65,66^, yet stromal cells showed conserved downregulation of collagen, fibronectin, and contractile genes. Because ECM proteins can accumulate through reduced turnover, declining transcription does not preclude accumulation of the age-associated stiffer matrix environment. Lower transcript levels may instead reflect changes in stromal cell states – or a compensatory response to changes in the ECM environment of the ovarian tissue.

Pathway-level, module-level, and trajectory analyses converged on granulosa cells as the most involved/affected cell type in the context of ovarian aging. GO terms related to autophagy, lysosome, and inflammatory response increased with age, whereas DNA repair and chromatin organization terms decreased. Among human-derived WGCNA modules, a module enriched for cell division and chromosome segregation was repressed with age, and a metabolic module was the most conserved, being depleted with age in six of seven mouse datasets. In pseudotime analysis, granulosa cells from older individuals occupied later pseudotime positions in both species, but follicle-stage composition shifted differently: the atretic fraction increased in human, whereas the preantral fraction declined in mouse. Together, these results suggest that aging moves granulosa cells toward growth-restrained, stress-responsive states, while the follicle-stage context in which this occurs differs between species.

In stromal cells, translational and mitochondrial terms increased with age, whereas cytoskeletal and adhesion terms decreased. In theca cells, a human module enriched for oxidative phosphorylation and translation was repressed with age but was not consistently recapitulated in mouse. These results indicate that energy and protein synthesis programs are remodeled in a cell-type-specific and species-dependent manner.

Leveraging the power of a multi-species, multi-dataset meta-analysis, we were able to identify 20 cross-species significant ASGARD-nominated compounds predicted to counteract age-associated transcriptional signatures. As an important sanity check, several of the ASGARD-targeted pathways are directly connected to the core ovarian aging signatures identified in this study. Dexamethasone had the broadest cross-species support, reaching significance in human, macaque, and mouse. As a glucocorticoid receptor agonist, it may temper the inflammatory programs elevated in the aging ovary, and, more importantly, aging granulosa cells ^67^. Clinical data in non-aging contexts support its plausibility as an ovarian modulator: in letrozole-resistant PCOS, adding low-dose dexamethasone increased ovulation, and in a single-blind IVF/ICSI trial in women with PCOS, adjuvant dexamethasone was associated with higher clinical pregnancy rates and lower gonadotropin requirements ^68,69^. However, future studies (outside of the scope of this meta-analysis) will need to establish its impact in the context of the aging ovary as a candidate intervention.

Two dihydropyridine L-type calcium channel blockers also ranked highly: nitrendipine, significant in human, mouse, and goat, and nicardipine, the top-ranked compound, supported by mouse and macaque ^70,71^. This class-level recurrence strengthens the signal beyond any single compound and points to calcium-dependent signaling, which is central to follicular physiology. Vorinostat, a histone deacetylase inhibitor ^72^, aligns with the reduced chromatin and genome maintenance programs in aged granulosa cells, and parthenolide, which inhibits NF-κB signaling through IκB kinase ^73^, aligns with the elevated inflammatory response terms.

The mTOR inhibitors sirolimus (rapamycin) and temsirolimus ^74,75^ were among the next-ranked compounds, supported mainly by mouse datasets. Indeed, in mice, short-term rapamycin treatment preserved the follicle reserve and extended ovarian lifespan ^76,77^. Importantly, rapamycin is currently the most clinically advanced candidate for ovarian aging – with human trials underway. Indeed, the ongoing VIBRANT pilot trial (NCT05836025) is testing low-dose weekly rapamycin in women aged 38–45 years, with ovarian reserve markers as outcomes, although peer-reviewed results have not yet been reported. Separately, daily low-dose rapamycin before IVF in 100 women increased embryo yield and clinical pregnancy rates, and the same study reported increased ribosome-related processes in aged cumulus cells ^78^. Given the elevation of genesets related to protein translation control we observed in the aging stroma, whether mTOR inhibition acts on these programs *in vivo* warrants further study. Because glucocorticoid, calcium, chromatin, and mTOR pathways all regulate steroidogenesis and ovulation, any effects of these compounds are expected to be dose-dependent and context-dependent. At this stage, these compounds remain candidates for mechanistic testing rather than therapeutic recommendations.

Finally, it is important to highlight that our findings are still limited primarily by the scope of available data. Human ovaries are large and regionally heterogeneous, and sections profiled by single-cell or single-nucleus RNA-seq do not – by design – reflect whole-organ composition ^79^. Public datasets are modest in size/biological replication, and remain uneven across species, platforms, and age windows, which unfortunately reduces statistical power and may bias estimates of direction in any given species. This is especially of note for the macaque and goat species, each represented by a single dataset, the goat dataset with only one replicate per age group. In contrast – likely due to ease of access and expertise – mouse datasets contributed the most samples, age coverage and therefore the most power – particularly in the ASGARD drug repurposing meta-analysis. We attempted to mitigate this effect by summarizing at the species level before cross-species integration, but it remains an important caveat. Another caveat to note is that ASGARD predictions rely on LINCS L1000 reference profiles, which were largely derived from cancer cell lines, and may not provide a perfect view of the impact of the drug in healthy, primary cells of different identity/state. Finally, since most of the available data was droplet-based and scRNA-seq rarely captures oocytes, our analysis has a blind spot for inference on oocyte-intrinsic aging pathways and whether they are concordant or divergent from the somatic compartment of the aging ovary.

Future work should include region-aware sampling of human ovaries; larger, stage-matched, species-balanced cohorts spanning pre-, peri-, and post-menopausal (or estropausal) stages; and orthogonal modalities such as spatial transcriptomics, sc/snATAC-seq, and proteomics. Mechanistic studies should test whether modulating FSH signaling, DNA repair, or cell-cycle restraint shifts granulosa cell states, and candidate compounds, including mTOR inhibitors and dihydropyridines, should be evaluated in primary cell culture, preclinical animal testing, and explant or organoid models. As more datasets accumulate, the same workflow can be reapplied to produce higher-confidence estimates at finer cell-state resolution and provide an ever-improving framework to understand (and ultimately mitigate) the pathways driving ovarian aging.

## Methods

### Datasets

We curated publicly available ovarian single-cell and single-nucleus RNA-seq datasets from sequencing repositories. Our inclusion criteria for the meta-analysis were (i) ovarian tissue origin with explicit sample labels, and (ii) separable age groups (no pooled or mixed-age libraries).

Raw FASTQ files were reprocessed for all datasets, except for the Tabula Sapiens, where a processed h5ad object was obtained due to data access restrictions (see below). All datasets underwent a consistent QC workflow, followed by harmonized annotation (see below).

For humans, we included three datasets spanning early adulthood to post-menopause: GSE202601 (snRNA-seq; 23–54 years) ^14^, GSE255690 (23, 38, and 48 years) ^15^, and the ovary subset of Tabula Sapiens (TS; 26, 55, and 56 years) ^22^. TS was obtained as a processed object from CZ CELLxGENE (https://cellxgene.cziscience.com/datasets). Ovarian cells were extracted and subjected to the same downstream QC thresholds, normalization, and annotation as the other datasets, without realignment.

For cynomolgus macaque (*Macaca fascicularis*), we analyzed GSE130664 generated by STRT-seq (hand-picked, sorted cells; 5 *vs*. 19 years) ^16^; because sampling fractions cannot reflect true tissue composition (but can still accurately represent transcriptomes of profiled cells), this dataset was excluded from proportion analyses, but used for transcriptional comparisons.

For mouse (*Mus musculus*), we included seven cohorts from five datasets/accessions, all generated on the 10x Genomics platform. Natural aging cohorts comprised GSE290742 (2 vs 9 months), GSE232309 (3 vs 9 months), E-MTAB-12889 (9, 12, and 15 months), E-MTAB-11491 (4 vs 18 months), and PRJNA863443-AC (4 vs 20 months). We also included control animals from two intervention studies within PRJNA863443: *Foxl2*^⁺/⁺^ animals from a *Foxl2* haploinsufficiency model (F+; 4, 9, and 17 months) and vehicle-injected controls from a VCD-induced ovarian failure model (Veh; 4, 6, 11, and 13 months).

For goat (*Capra hircus*), we included PRJNA1010653, generated on the 10x Genomics platform (2 vs 10 years). This dataset included one sample per age group, so its results were interpreted mainly in terms of consistency with the other species.

Detailed information on the datasets is provided in Table S1.

### Data processing

For 10x Genomics datasets, raw sequencing reads were processed with Cell Ranger (v. 7.1.0, 10x Genomics) ^80^. Reads were aligned to species-specific reference genomes provided by 10x Genomics or Ensembl: GRCh38-2020-A for human and mm10-2020-A for mouse. For goat, (*Capra hircus*) a custom 10x Genomics-compatible reference was constructed with Cell Ranger using the cellranger mkref command and the NCBI RefSeq assembly GCF_001704415.2 fasta and gtf.

To minimize the impact of contamination from ambient RNA, Cell Ranger expression matrices were corrected using DecontX from the celda package (v. 1.14.2) ^81^ in R (v. 4.4.1), with both raw and filtered feature-barcode matrices used to estimate ambient RNA levels. Doublets were removed through a two-step strategy. First, DoubletFinder (v. 2.0.3) ^82^ was run on each sample independently, with expected multiplet rates provided by 10x Genomics and the optimal pK parameter identified via parameter sweep. Predicted doublets were flagged and excluded. In parallel, scds (v. 1.10.0) ^83^ was applied using the cxds_bcds_hybrid function, and cells identified as doublets by either approach were discarded from downstream analysis.

The resulting filtered single-cell expression matrices were analyzed in Seurat (v. 4.3.0) ^84^. For Tabula Sapiens dataset, the processed count matrix entered the pipeline at this step. Initial quality control removed cells with fewer than 500 detected features, total RNA counts outside 500–100,000, mitochondrial RNA content greater than 15%, or DecontX contamination scores above 0.25. After quality control, datasets were normalized with SCTransform, regressing out the number of detected features, mitochondrial percentage and DecontX contamination score. The number of QC cells in each dataset is reported in Table S1.

For the STRT-seq dataset, raw sequencing reads were first trimmed to remove template switch oligo (TSO) and poly(A) tail sequences using Cutadapt (v. 4.4) ^85^. Reads containing more than 10% low-quality bases (N) or adapter contamination were discarded. Cleaned reads were then aligned to the *Macaca fascicularis* reference genome (Ensembl Macaca_fascicularis_5.0, Ensembl release 115) using kallisto-bustools (v. 0.27.3) ^86^. Barcode error correction was performed using a whitelist of valid barcodes provided by the original study ^16^. The resulting single-cell expression matrices were analyzed in Seurat (v. 4.3.0) ^84^. Cells were retained if they passed three QC thresholds: more than 500 detected features, more than 500 unique molecular identifiers (UMIs), and mitochondrial RNA content below 15%. After quality control, the dataset was processed following the same analysis pipeline used for the 10x Genomics datasets. The number of cells post-processing in each dataset is reported in Table S1.

### Cell type annotation

For the 10x Genomics datasets, cell type annotation was performed using a multi-step strategy that combined automated reference-based approaches with manual marker-based validation. First, *Ptprc*^⁺+^ and *Ptprc*⁻ cell populations were identified using scGate ^87^. Cells expressing *Ptprc* (encoding the Cd45 pan-immune cell surface protein) were classified as immune, while those lacking *Ptprc* expression were labeled as non-immune.

The two subsets were then annotated independently for greater granularity and accuracy. For immune cells, SingleR (v. 1.8.1) ^88^ was applied using datasets from celldex (v. 1.4.0) as a transcriptomic reference ^89^, and the resulting annotations were compared with the output of marker-based scSorter (v. 0.0.2) ^90^ and scType (v. 1.0) ^91^. Markers used for each cell type are listed in Table S2. For non-immune cells, only marker-based scSorter and scType were used, due to the limited availability of high-quality ovarian single-cell reference datasets; annotations were supplemented by manual curation based on canonical ovarian marker expression (Table S2). Final cell type labels were assigned using a majority voting scheme across computational outputs, together with a final manual annotation. All analyses were conducted using Seurat (v. 4.3.0) framework ^84^.

For the STRT-seq dataset, cell type annotation was based on the classifications provided by the original study ^16^, as cells had been manually selected prior to sequencing. These reported labels were retained for downstream analyses to ensure consistency with the experimental design.

### Data integration

To account for batch effects within the human and mouse datasets, integration was performed on the independent preprocessed datasets for each species using Seurat (v. 4.3.0) ^84^. First, Seurat objects for each species were merged. Principal component analysis (PCA) was then performed on the merged datasets, and batch effects were assessed by visualizing PCA embeddings colored by dataset identity. Subsequently, batch correction was applied via Seurat’s anchor-based reciprocal PCA (RPCA) approach. The corrected embeddings were then used for any downstream joint analysis (*e.g*. UMAP visualization).

### Cell-type proportion and Augur analysis

Cell-type proportions were evaluated using scProportionTest (v. 0.0.0.9000) in R ^33^, which applies a permutation-based framework to compare cell-type abundances between groups. This method estimates relative changes in proportions and provides confidence intervals for each cell type. Analyses were performed at two levels: the broad division of *Ptprc*⁺ versus *Ptprc*⁻ populations, and the finer resolution of individual cell subtypes within each group. Samples were assigned to reproductive phases based on age. Human donors were classified as pre-menopausal (<45 years), peri-menopausal (45–51 years), or post-menopausal (≥52 years). Mice were classified as pre-estropausal (<9 months), peri-estropausal (9–15 months), or post-estropausal (>15 months). For datasets spanning two reproductive phases, samples within each phase were pooled and the two phases were compared (e.g., pre-*vs*. peri-menopause for GSE255690). For GSE202601, which spanned all three phases, and EMTAB12889, which was confined to peri-estropause, the youngest and oldest ages were compared directly. For PRJNA863443-F+, 4- and 9-month samples were compared to ensure largest available replication within this dataset, as the 17-month group comprised a single sample (n = 1). Goat samples were compared as young (2 years) vs. aged (10 years).

To identify cell types whose transcriptome was most affected by aging, we applied Augur (v. 1.0.3) ^29^. Analyses were conducted both pairwise and across all groups simultaneously, depending on dataset design. For datasets with two age groups, comparisons were made directly between young and old subjects, while for datasets with three ages, each age was treated as a distinct group.

### Pseudobulk analysis for differential gene expression

Pseudobulk differential gene expression analysis was performed for each dataset. Single-cell transcriptomic data were aggregated at the cell type level using muscat (v. 1.18.0) ^92^ to generate pseudobulk expression profiles. Cell types were retained if they contained at least 25 cells per sample in at least 75% of samples within a dataset.

Differential gene expression was then assessed using DESeq2 (v. 1.44.0) ^38^, applying a linear model with age as the main variable. To account for hidden sources of variation, surrogate variables were estimated with sva (v. 3.52.0) ^93^, and, when necessary, surrogate variable correction was applied using the removeBatchEffect() function from limma (v. 3.60.6) ^94^. Variance-stabilized counts were generated with getVarianceStabilizedData(), and differentially expressed genes (FDR < 0.05) were visualized using bubble plots.

### Humanization of DESeq2 results

Ortholog mapping was performed in R using biomaRt (v.2.60.1; Ensembl 115) ^95^. Species-specific Ensembl marts (*Macaca fascicularis, Mus musculus, Capra hircus*) were queried for ensembl_gene_id, external_gene_name, hsapiens_homolog_ensembl_gene, hsapiens_homolog_associated_gene_name, and hsapiens_homolog_orthology_type. Returned mappings were filtered to one-to-one orthologs (ortholog_one2one) with non-empty human symbols and deduplicated by the source gene name. For each dataset and cell type, DESeq2 result tables were standardized to data frames, restricted to mapped genes, and row names were replaced with the corresponding human symbols.

### Consistent DEG analysis

Humanized, cell-type–resolved DESeq2 results were used to identify differentially expressed genes (DEGs) with consistent age-associated changes across species. For species represented by multiple datasets (human, mouse), per-dataset p-values were combined within species using the inverse-normal method in metaRNASeq (v. 1.0.8; invnorm) ^39^, followed by Benjamini–Hochberg correction to obtain species-level FDR values. For single-dataset species (macaque, goat), DESeq2 FDR values were used directly.

Within each species, genes with FDR < 0.10 were considered significant and were separated by direction of change. For granulosa and stromal cells, conserved DEGs were defined as genes significant in at least three of the four species with a consistent direction of change. Theca cells were not detected in the macaque dataset, so conserved theca DEGs had to be significant, with a consistent direction, in all three remaining species (human, mouse, and goat). For species overlaps, significant sets were defined as FDR < 0.10 per species, and counts-only Venn diagrams (human, macaque, mouse, goat) were produced using ggVennDiagram (v. 1.5.4) ^96^.

### Gene Ontology analysis using mitch

Humanized, cell-type–resolved DESeq2 results were subjected to Gene Ontology (GO) enrichment with the mitch framework (v. 1.16.1) ^59^ in R. GO gene sets were assembled from Ensembl BioMart using the Human genes (GRCh38.p14) dataset. At export, filters excluded pseudogene and non-coding RNA gene types, and GO annotations carrying NAS or TAS evidence codes (i.e. non-experimental author statements) were removed. The tab-delimited file was imported into R, trimmed and deduplicated to retain unique gene–term pairs, and used to construct GO gene set definitions.

For each cell type, DESeq2 result tables from available datasets were standardized, merged by gene symbol, and imported into mitch (mitch_import, DEtype=“DESeq2”). Enrichment was then computed with mitch_calc using the custom GO and Reactome sets (minimum set size = 10; priority = “significance”). The global multivariate statistic across contrasts was corrected across sets by Benjamini–Hochberg; significant terms were defined as p.adjustMANOVA < 0.05. Contrast-specific signed effect sizes returned by mitch were visualized as heatmaps using ComplexHeatmap (v. 2.21.1) ^97^.

### WGCNA and cross-species enrichment analysis

Weighted gene co-expression network analysis (WGCNA) ^30^ was performed on human, cell-type–resolved variance-stabilized counts from DESeq2 to identify age-associated modules using WGCNA (v. 1.73). Analyses were restricted to the intersection of genes present in all contributing datasets. The soft-thresholding power was chosen separately in each dataset with pickSoftThreshold. Consensus modules were identified with blockwiseConsensusModules (corType=“bicor”, deepSplit=2, mergeCutHeight=0.20, maxBlockSize=40000, and minimum module size ≈2.5% of genes). Modules were declared age-associated if (i) the ME–age effects were concordant in direction across datasets and (ii) the meta-analytic Benjamini–Hochberg FDR < 0.1.

Human-derived module gene sets were then tested for consistency across datasets and species by gene set enrichment analysis (GSEA) via clusterProfiler (v4.12.6) ^98^ against age-ranked gene lists from each human, macaque, mouse, and goat dataset; significance was set at FDR < 0.10. Functional characterization of granulosa modules used over-representation analysis (ORA) via clusterProfiler (v. 4.12.6) ^98^ with org.Hs.eg.db (v. 3.19.1), reporting GO terms at FDR < 0.05.

### Pseudotime analysis of granulosa cells

Granulosa cells from human and mouse datasets were integrated separately for each species. GSE255690 granulosa cells were retained here because cell-level analyses did not require the minimum cell counts applied for pseudobulk aggregation. Trajectories were inferred with Monocle3 (v. 1.4.27) ^31^ using cluster_cells and learn_graph, with root nodes selected manually (order_cells) and verified by a positive correlation between pseudotime and age.

To avoid pseudoreplication, pseudotime was averaged per donor or animal (human, n = 14; mouse, n = 52). Differences across age groups (human: 23–29, 38–49, and 51–56 years; mouse: 2–6, 9–15, and 17–20 months) were tested with the Kruskal-Wallis test, followed by pairwise Wilcoxon rank-sum tests with Benjamini-Hochberg correction. Follicle stages (preantral, antral, atretic, and other) were assigned using published marker-based schemes ^34^ and summarized by age group. Over-representation analysis of increasing and decreasing genes combined was performed with clusterProfiler (v. 4.12.6) ^99^ against GO Biological Process terms, and terms with FDR < 0.05 were considered significant.

### ASGARD drug repurposing and cross-species meta-analysis

Humanized, cell type–resolved differential signatures (limma on log-normalized counts; case = old, control = young; minimum 3 cells per cell type) were generated for each dataset using ASGARD (v. 1.0.0) ^32^. Each cell-type signature was queried with ASGARD against LINCS L1000 reference profiles. ASGARD produced drug-level statistics for each cell type, which were aggregated across cell types within a dataset using a weighted scheme (cell type frequency × signal strength) to give a Drug.therapeutic.score and a combined p-value per drug. For species with multiple datasets, drug-level p-values were combined across datasets using Fisher’s method to obtain a per-species p-value. Per-species p-values were then combined across the four species using Fisher’s method to obtain a cross-species combined p-value. Drugs were ranked by the combined p-value, and per-species p-values < 0.05 were considered significant.

### R shiny application generation

An interactive R shiny application of sc/snRNA-seq datasets across models was generated using “ShinyCell” (v.2.1.0) ^100^, and made available at https://minhooki.shinyapps.io/cross-species-ovary-scRNA/.

## Data availability

This study analyzed publicly available datasets. The human datasets are available from the Gene Expression Omnibus (GEO) under accession codes GSE202601 and GSE255690, and from Tabula Sapiens (https://cellxgene.cziscience.com/datasets). The cynomolgus macaque dataset is available from GEO under accession code GSE130664. The mouse datasets are available from GEO under accession codes GSE290742 and GSE232309, from the NCBI Sequence Read Archive under BioProject PRJNA863443, and from ArrayExpress under accession codes E-MTAB-12889 and E-MTAB-11491. The goat dataset is available from the NCBI Sequence Read Archive under BioProject PRJNA1010653. Details of each dataset, including sample ages and cell numbers after quality control, are provided in Supplementary Table S1.

## Code availability

The underlying code for this study is available in GitHub and can be accessed via this link https://github.com/BenayounLaboratory/Cross-species_ovarian_aging_analysis.

## Acknowledgements

This work was supported by USC Provost’s Undergraduate Research Fellowship to J.W.; Single-Cell Biology Data Insights award 2023-323351 from Chan Zuckerberg Initiative, Pew Biomedical Scholar award #00034120 from the Pew Charitable Trust, and generous gifts from Kathleen Gilmore, Dr. Eric Hennigan and Kayla Barnes to B.A.B.

Resources from Flaticon (www.flaticon.com) were used to generate some panels.

## Author contributions

M.K. and B.A.B. designed the study. M.K, J.W., X.L. and E.S performed computational analyses. M.K. and B.A.B. wrote the manuscript with input from all authors. All authors edited and commented on the manuscript.

## Competing interests

All authors declare no financial or non-financial competing interests.

## Supplementary figure legends

**Supplementary fig. 1 Marker gene profiles of *Ptprc^+^*cell types across datasets.** Marker gene dot plots of *Ptpr^+^*cell types from integrated human (**a**), macaque (**b**), integrated mouse (**c**) and goat (**d**) datasets.

**Supplementary fig. 2 UMAP plots of Augur AUC scores from individual datasets.**

**Supplementary fig. 3 Analysis of ovarian aging–associated differential gene expression in theca and stroma across datasets.** (**a** and **b**) Bubble plot of top consistent DEGs (FDR < 0.1) between human vs. macaque, human vs. mous) Venn diagram of overlapping DEGs (FDR < 0.1) across species for theca (**c**) and stroma (**d**).

**Supplementary fig. 4 Module trees of human WGCNA analysis.** (**a-c**) Module trees identified in granulosa – GSE202601 (**a**) theca – GSE255690 (**b**), and stroma (GSE255690 and GSE202601).

**Supplementary fig. 5 Association analysis of WGCNA modules and ovarian aging traits from human datasets.** (**a-c**) Heatmap of trait significance for WGCNA modules in granulosa (**a**), theca (**b**) and stromal cells (**c**).

**Supplementary fig. 6 GSEA of ovarian aging–associated WGCNA modules in theca and stroma across datasets.** (**a** and **b**) Cross-species GSEA enrichment (FDR < 0.1) of human theca (**a**) and stromal (**b**) cells-derived WGCNA module gene sets against macaque, mouse and goat datasets.

## Supplementary Tables

**Table S1. Summary of curated datasets.**

**Table S2. List of marker genes used for cell type annotation.**

## Notes

### Competing Interest Statement

The authors have declared no competing interest.

### Summary of Updates

- added additional datasets - updated analyses - updated interpretations

