## Supplementary Figures 1-12 for "Single-cell exploration of ovarian aging across mammalian models"

Supplementary figure 1

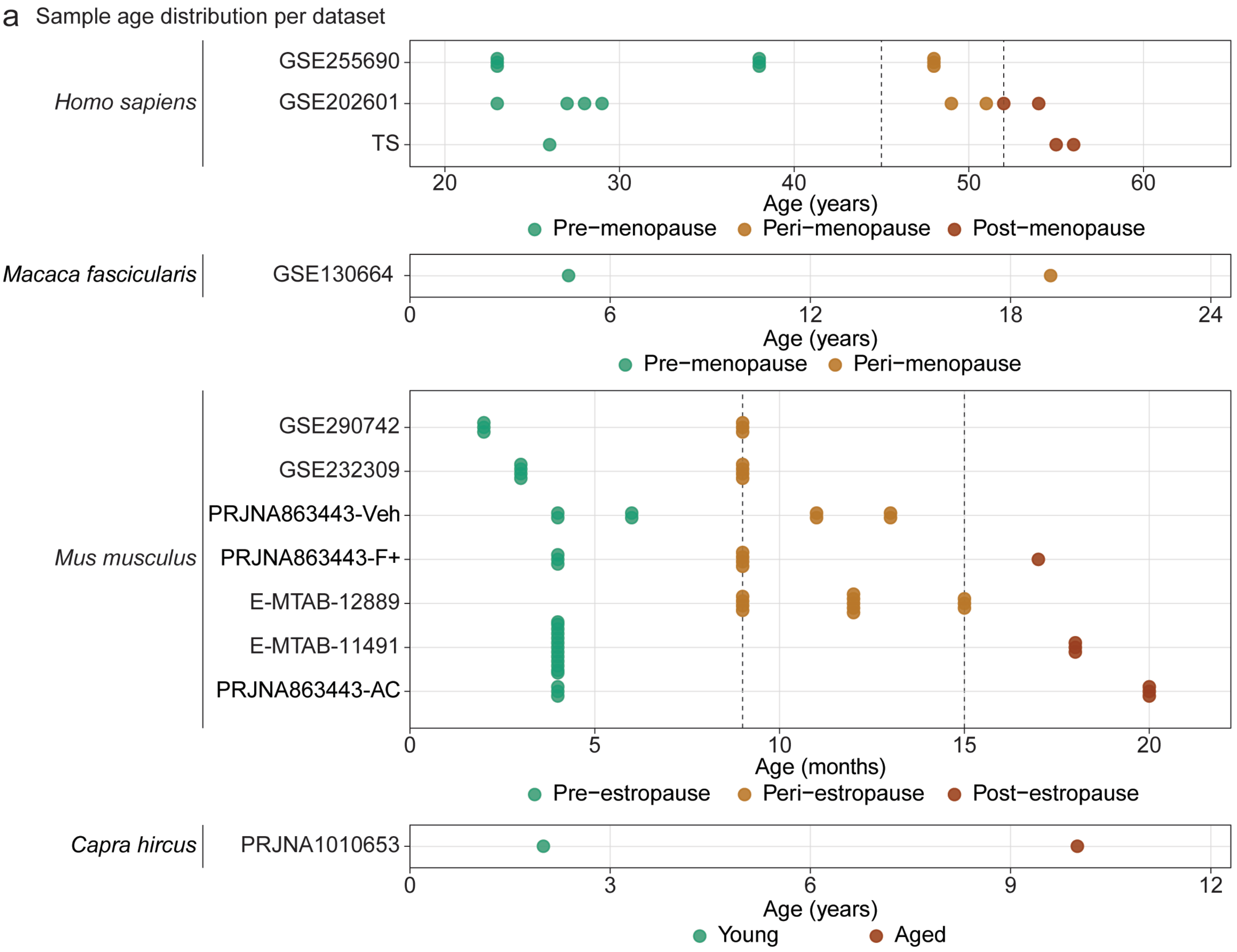

Supplementary figure 2

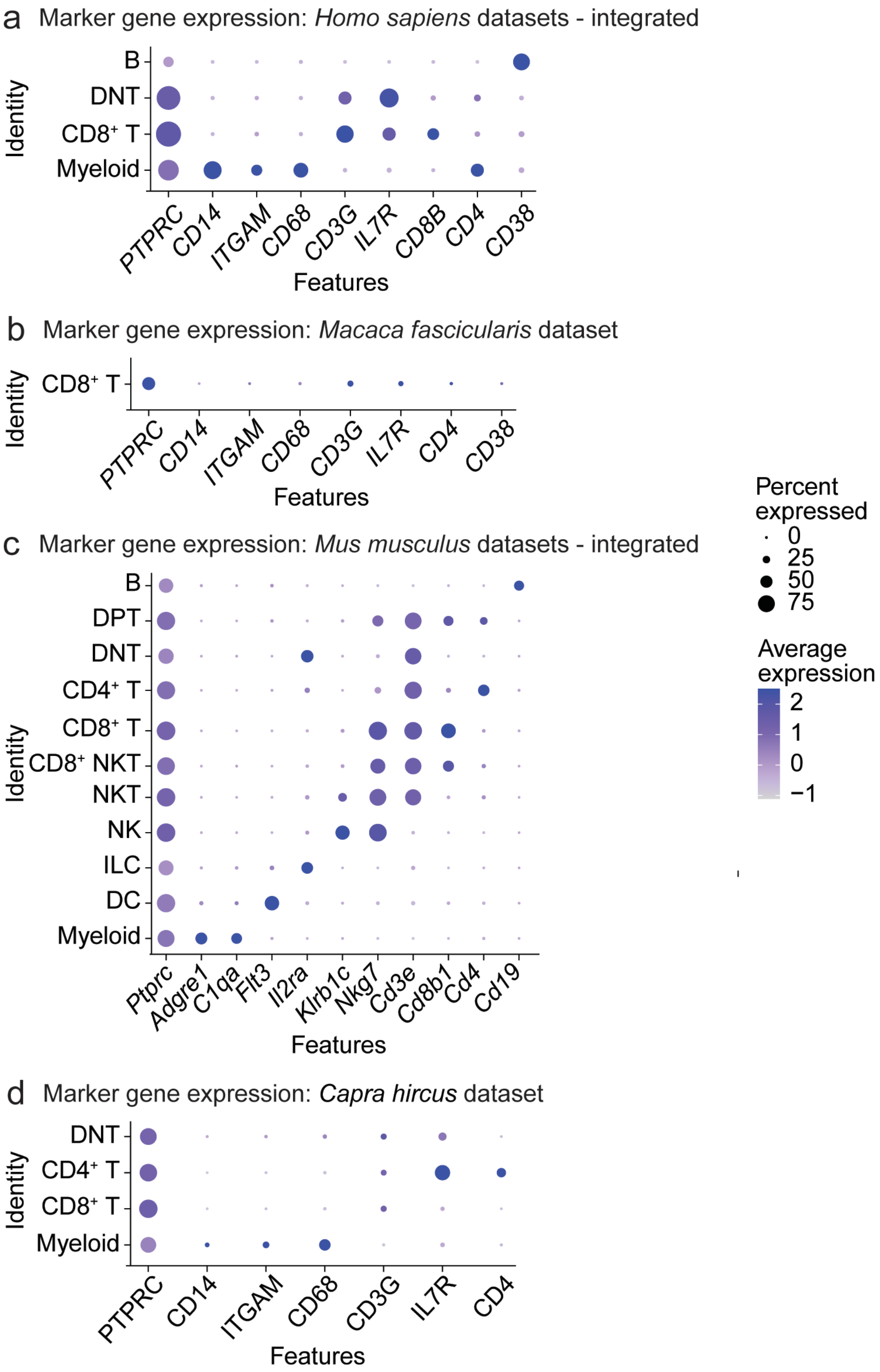

Supplementary figure 3

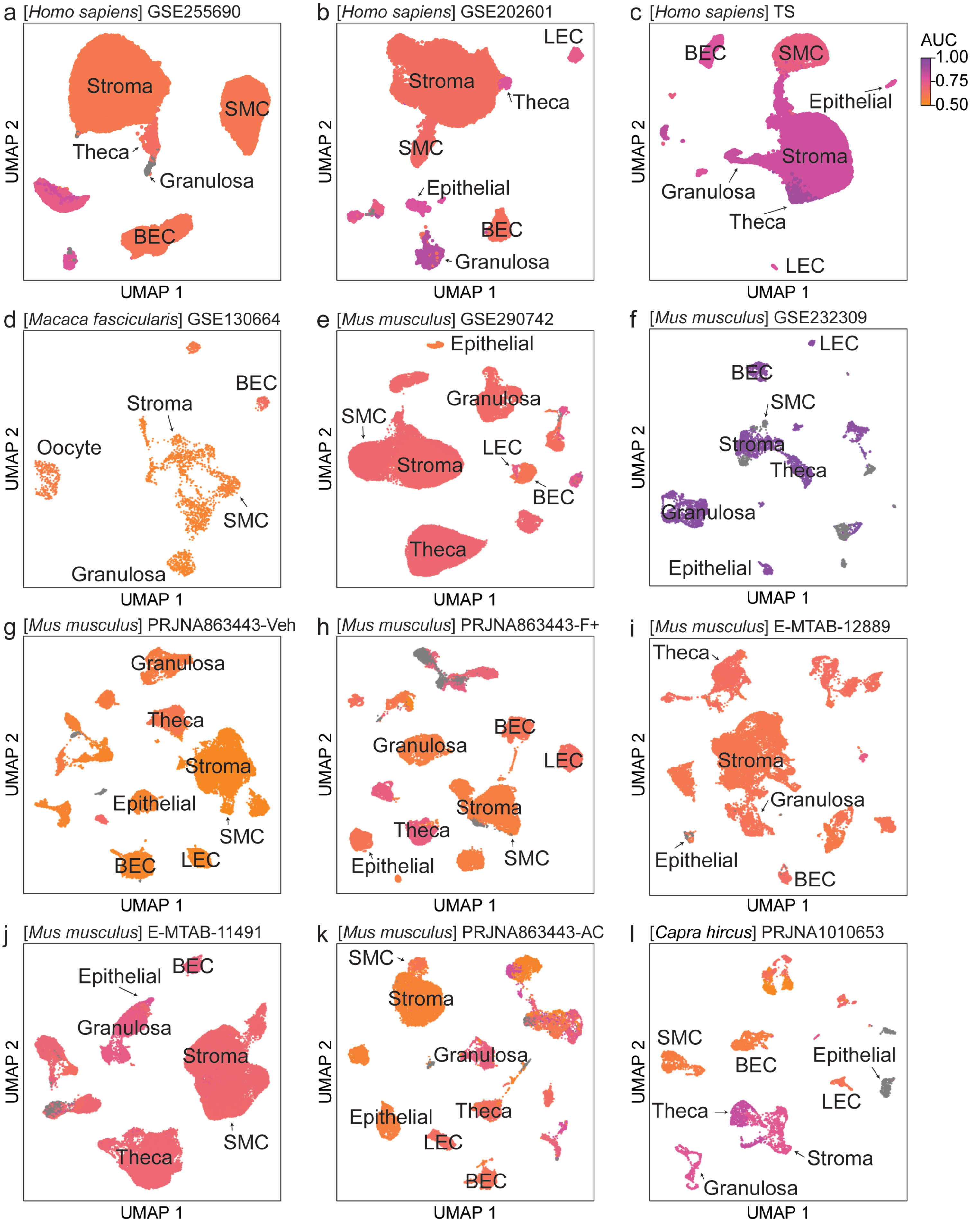

Supplementary figure 4

a Conserved DEGs (3 out of 4 species): Granulosa cells

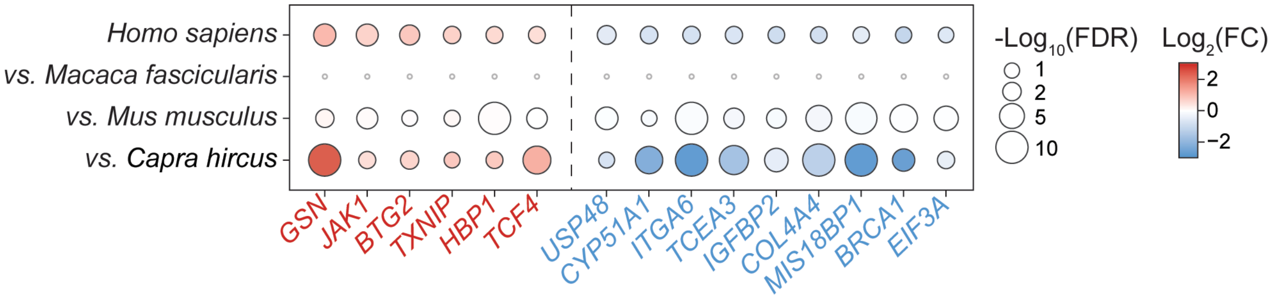

Supplementary figure 5

a Overlap of up-regulated DEGs: Theca cells (FDR < 0.1)

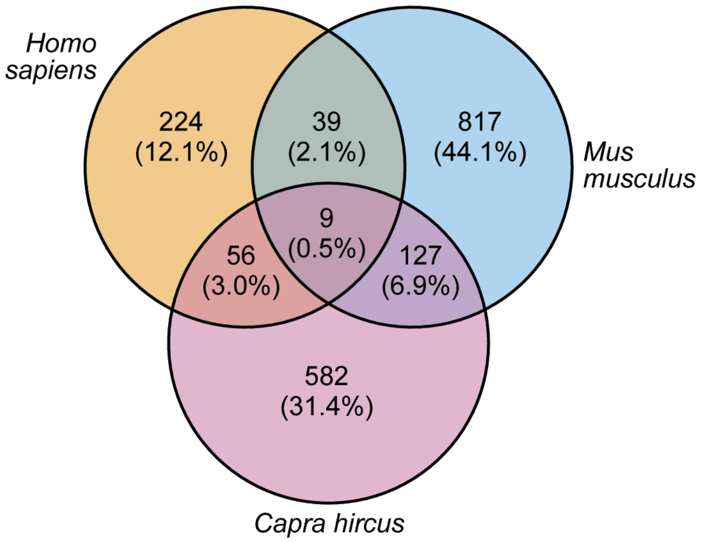


b Overlap of down-regulated DEGs: Theca cells (FDR < 0.1)

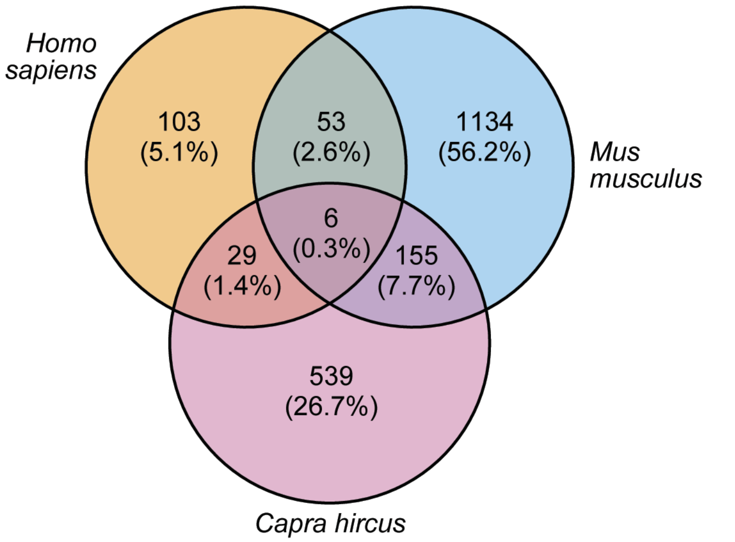


c Conserved DEGs: Theca cells

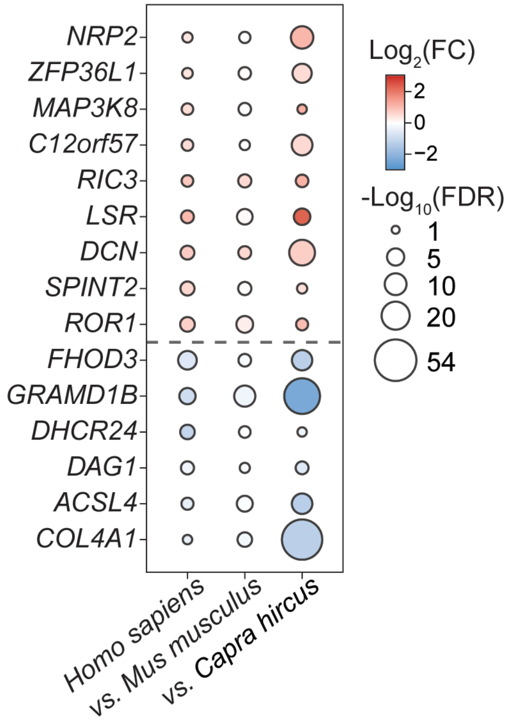

Supplementary figure 6

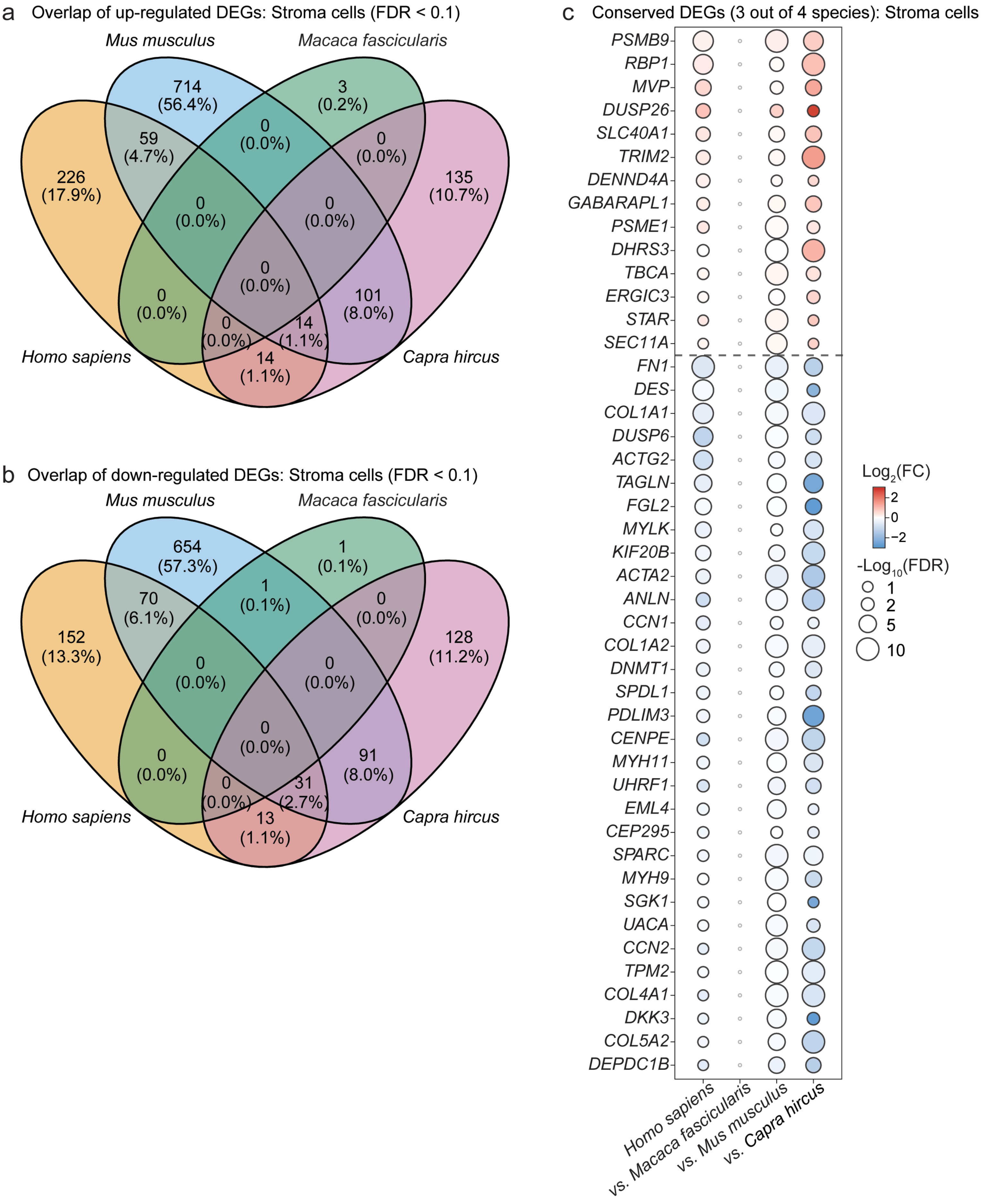

Supplementary figure 7

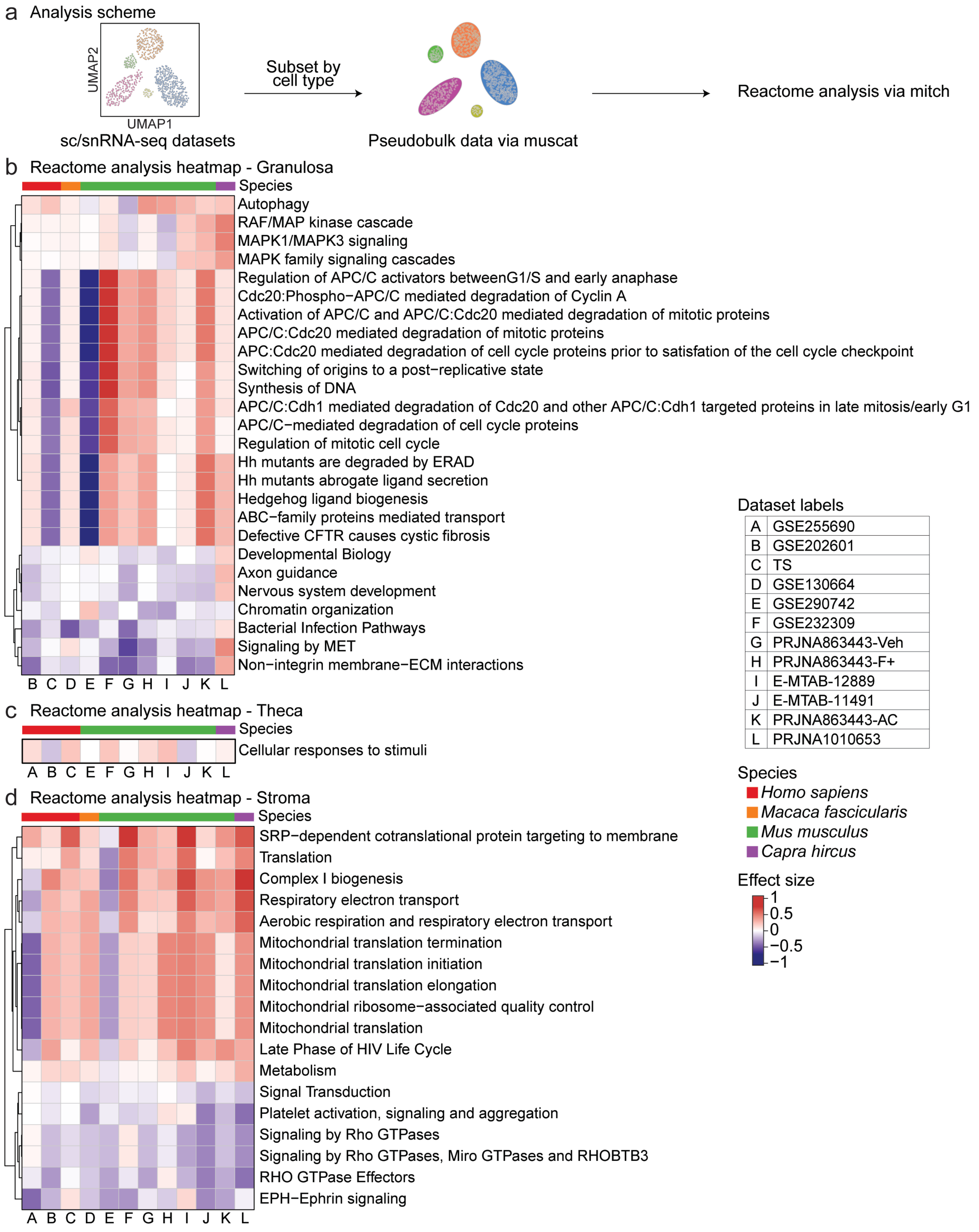

Supplementary figure 8

a Gene clustering dendrogram - Granulosa

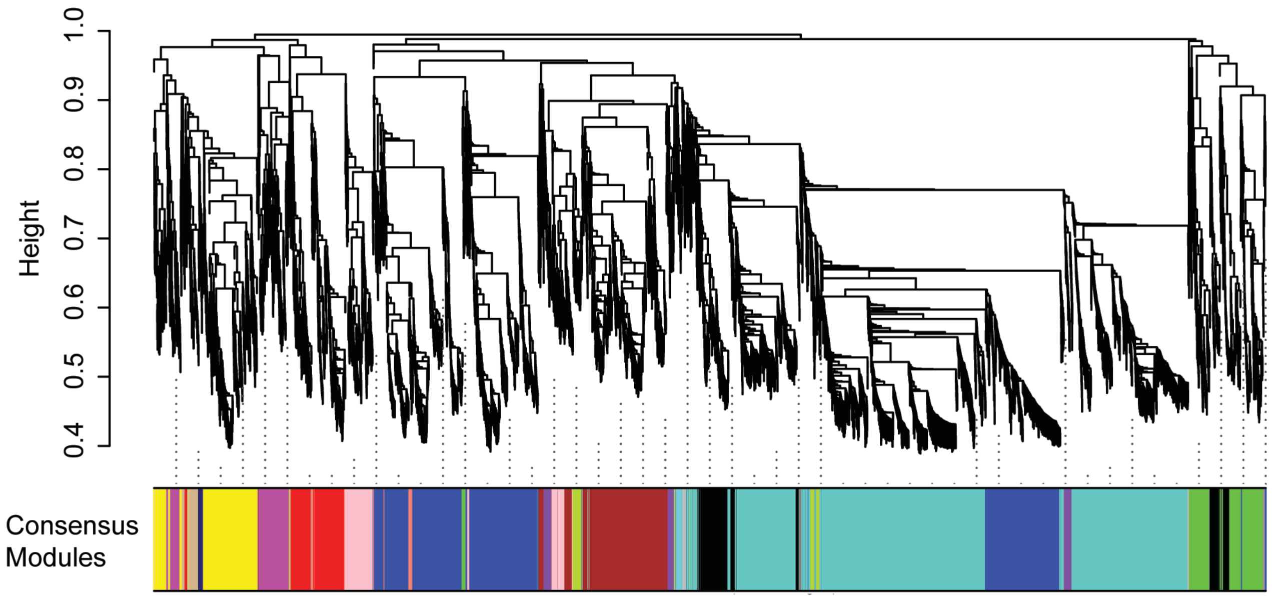


b WGCNA trait association - Granulosa

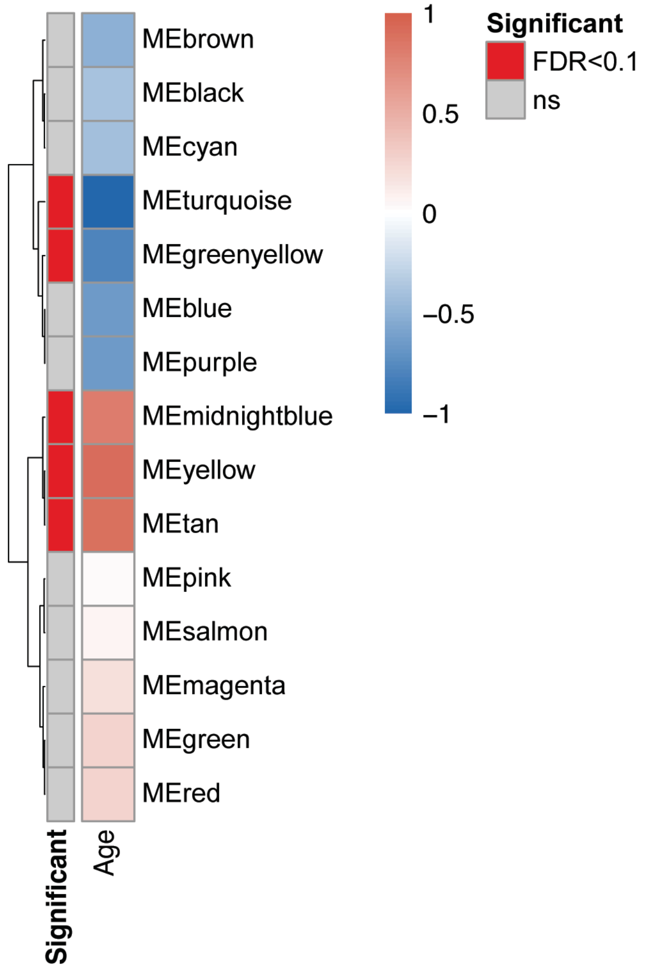

Supplementary figure 9

a Gene clustering dendrogram - Theca

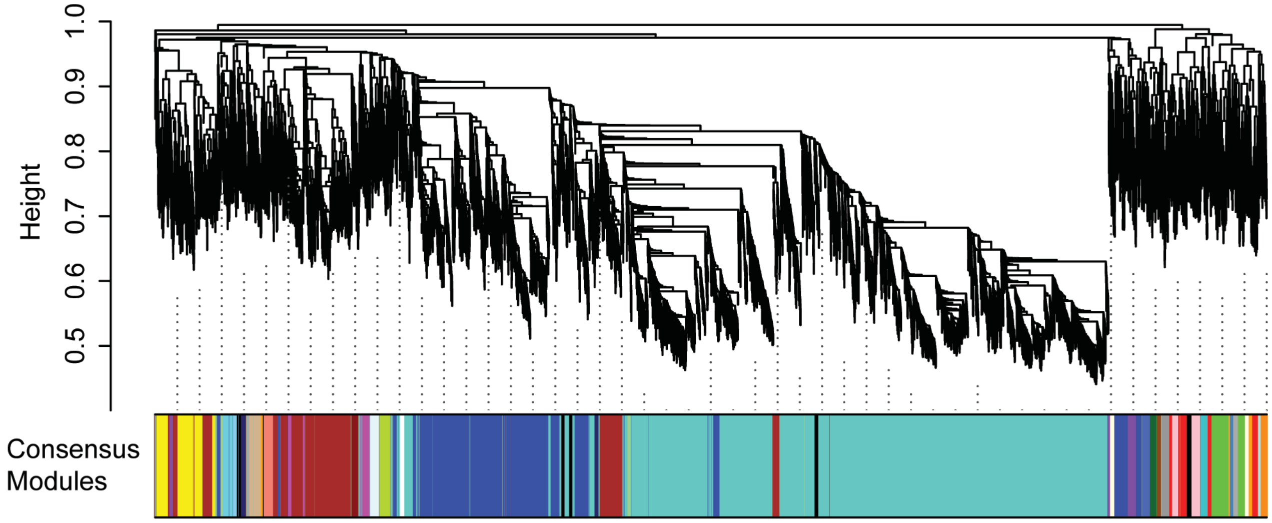


b WGCNA trait association - Theca

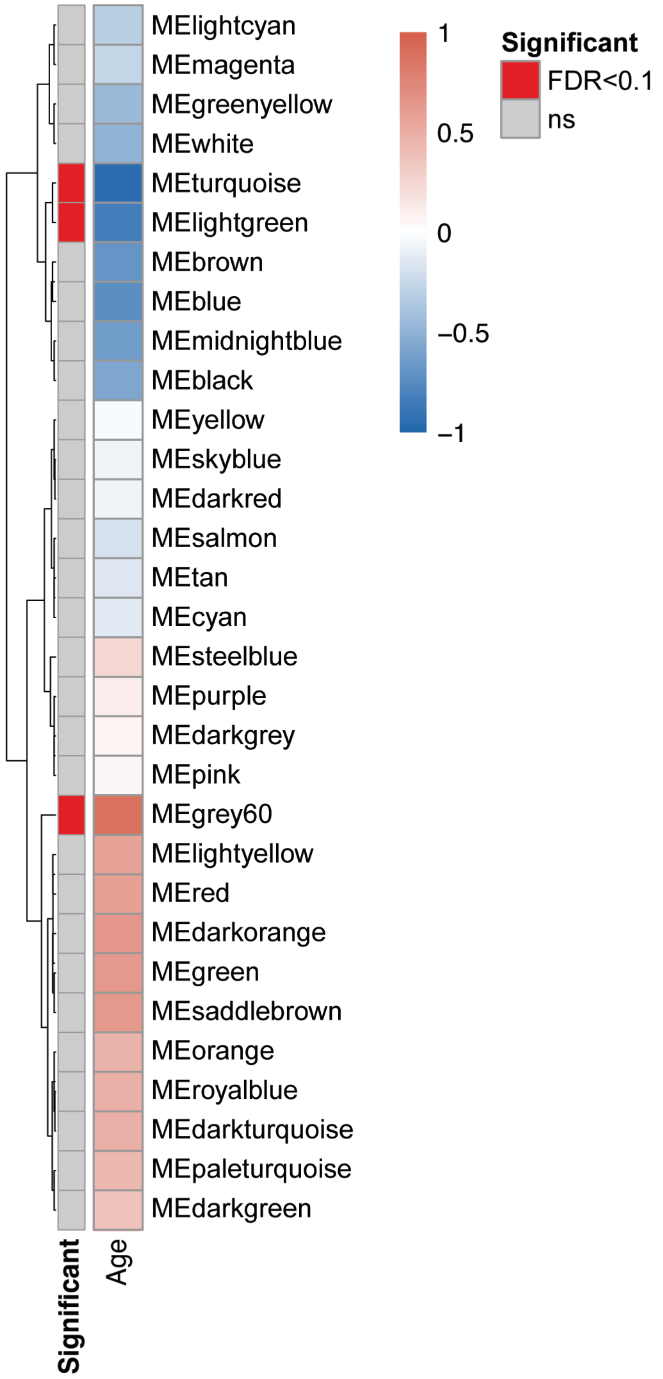

Supplementary figure 10

a GSEA: Theca WGCNA module gene sets

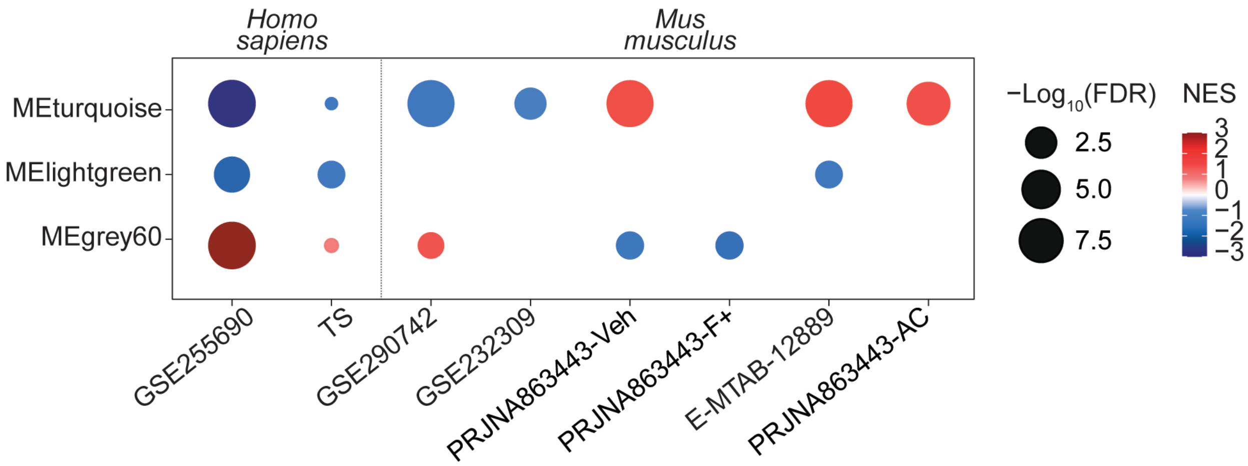


b Theca WGCNA module: MEturquoise

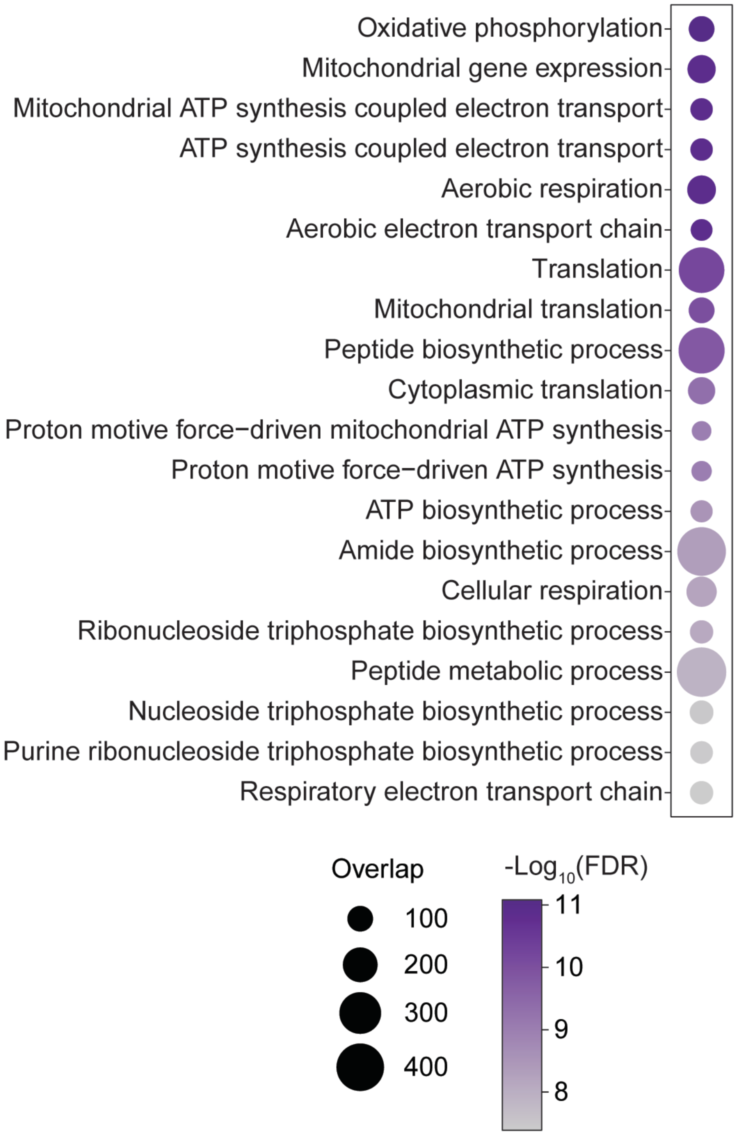


c Theca WGCNA module: MEgray60

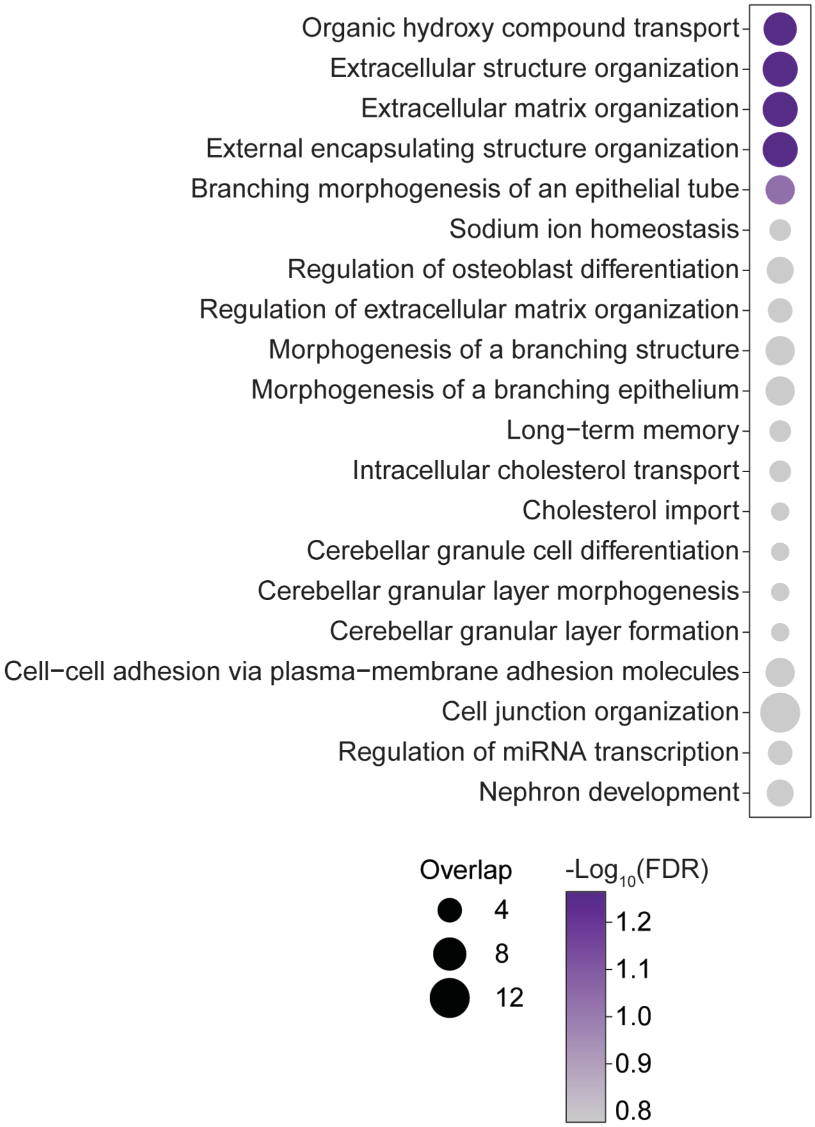

Supplementary figure 11

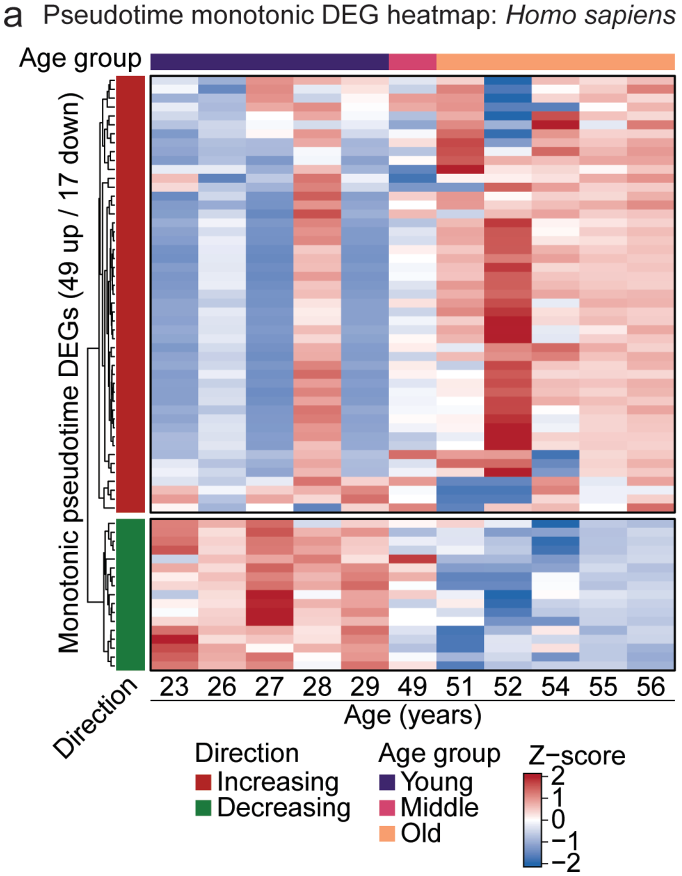


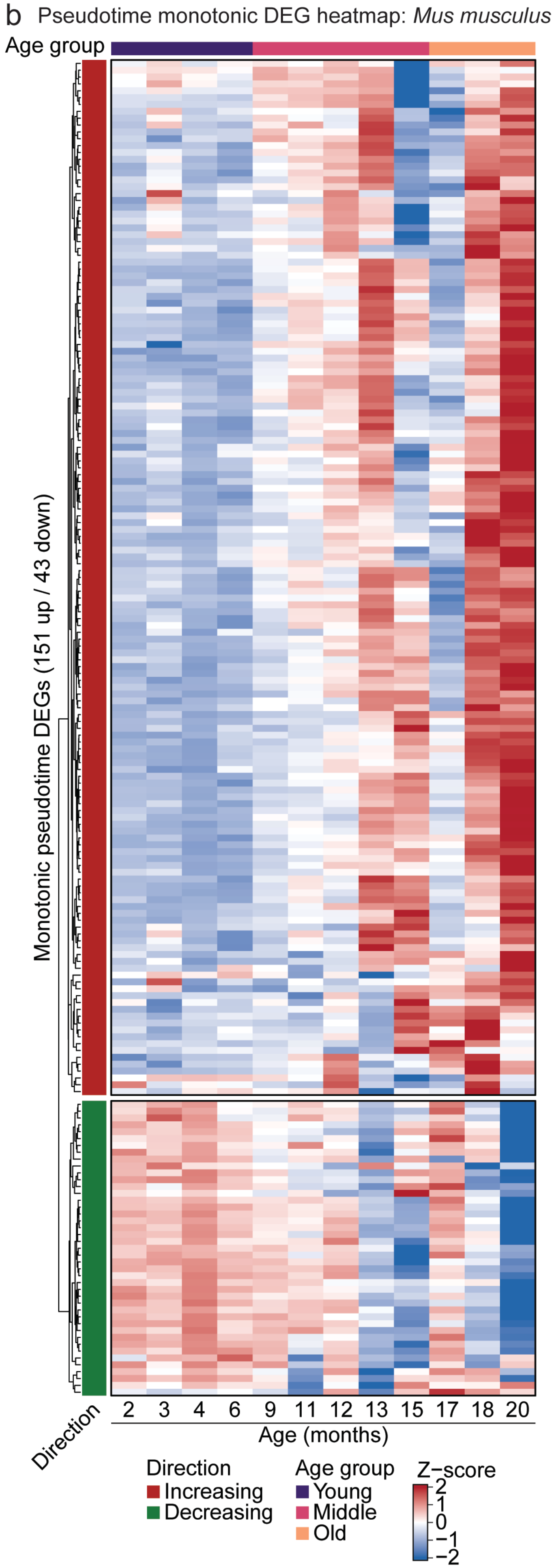

Supplementary figure 12

a Cross-species drug repurposing analysis

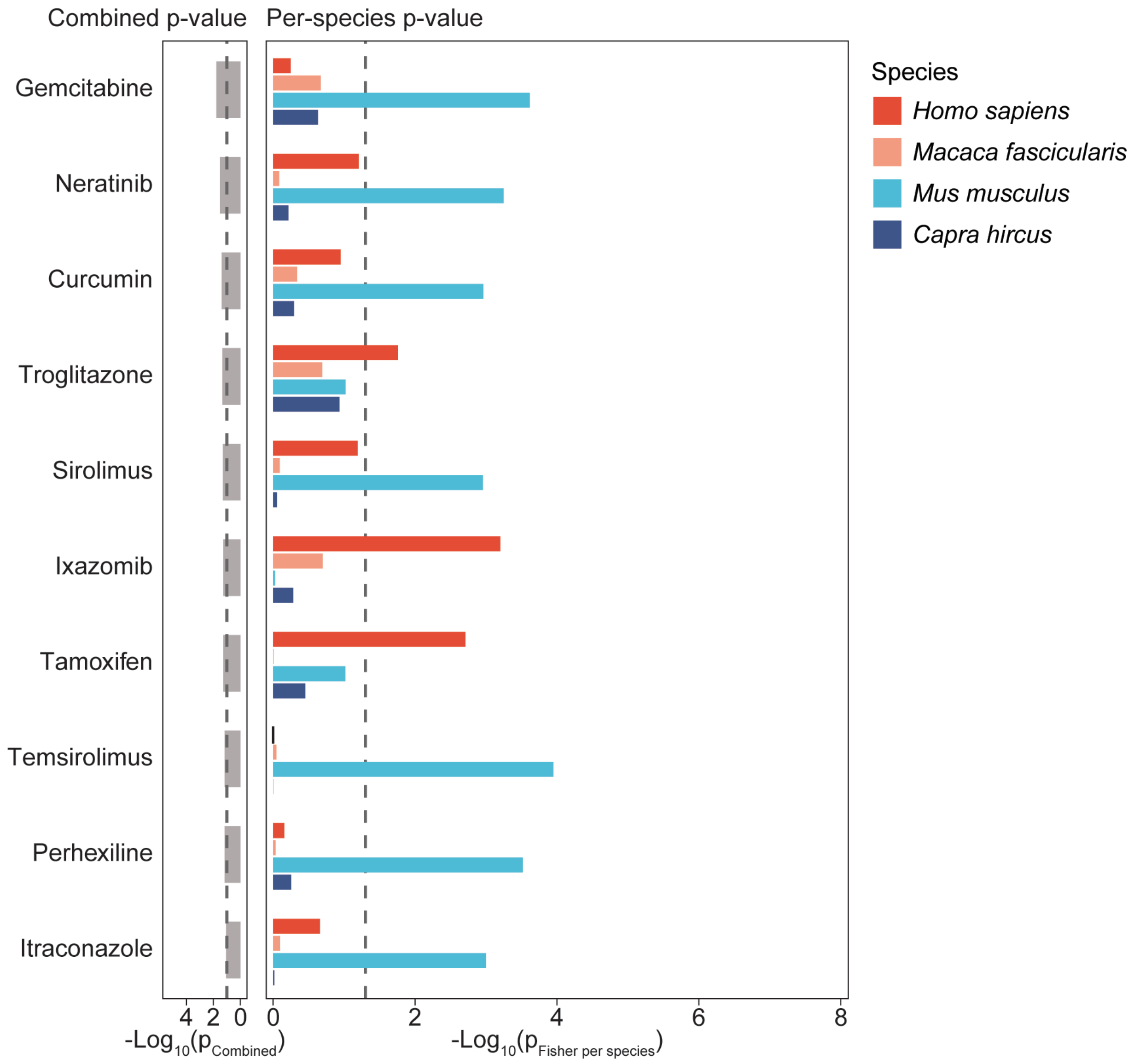
